# Prefrontal parvalbumin neurons as a target for enhancing cognition in non-pathological and 22q11.2 microdeletion syndrome mice

**DOI:** 10.1101/2025.03.31.645999

**Authors:** Tyler D. Dexter, Daniel Palmer, Ashley L. Schormans, Shahnaza Hamidullah, Meira M.F. Machado, Brian L. Allman, Timothy J. Bussey, Lisa M. Saksida

## Abstract

A failure of organized communication in the PFC is thought to contribute to the emergence of cognitive impairments in psychiatric diseases, with attentional deficits occurring as a fundamental symptom across various conditions. The 22q11.2 microdeletion syndrome is a rare genetic condition that confers a high risk for developing psychiatric and neurodevelopmental disorders, and mouse models have been shown to display attention impairments and PFC pathology that are relevant to clinical populations. Abnormalities in prefrontal parvalbumin-expressing neurons (PVNs) are part of the observed pathophysiology, and studies in rodents have shown that the direct manipulation of these cells can induce behavioral deficits that align with the cognitive symptoms observed in psychiatric diseases. In the present study, we expanded on the role of PVNs in supporting cognition by investigating their involvement in multiple aspects of attentional functions using a translationally relevant task of focused visual attention, in both non-pathological mice and a model of the 22q11.2 microdeletion syndrome. We observed that task-evoked prefrontal PVN activity was reduced in mice that exhibited poorer attention and in 22q11.2 mutant mice. While PVN activity was shaped across learning in non-pathological mice, mutant mice exhibited a lack of signal dynamics that coincided with attentional deficits. Importantly, we observed that task performance in both poor performing wild-types and 22q11.2 mutants could be alleviated by gamma frequency stimulation of PVNs. Thus, PVNs appear to be involved in the acquisition of task rules and execution of attention and continue to be a promising therapeutic target for cognitive dysfunction in disease.

## Introduction

Parvalbumin-expressing neurons (PVNs) are a component of canonical cortical circuitry and have been shown to be necessary for various prefrontal-cortex (PFC)-dependent cognitive processes such as working memory [1–4], cognitive flexibility [1, 4–9], and sustained attention [10, 11]. PVNs support cortical circuits by organizing the temporal responsiveness of local neurons and have been shown to facilitate the emergence of synchronous high-frequency gamma oscillations (30 – 100hz), which are associated with processing sensory information in the cortex [12–15]. Elevated prefrontal gamma activity is observed when rodents perform tasks that are sensitive to PVN disruptions, including working memory [16], task-switching [6, 7, 17, 18], and attention [10]. Cortical PVN dysfunction is strongly implicated in diseases that affect cognition, and have been particularly associated with psychiatric disorders such as schizophrenia [19–35], but also neurodegenerative diseases such as Alzheimer’s [36–40], Parkinson’s [41, 42], and Huntington’s Disease [43, 44]. Recently, a few studies have assessed the therapeutic potential of targeting prefrontal PVNs with low gamma stimulation to alleviate cognitive deficits in animal models of psychiatric disorders, and have shown that 30-40hz stimulation can improve impairments in cognitive domains such as sustained attention [11] and attentional set shifting [6, 9].

The 22q11.2 microdeletion syndrome (22qDS) is the result of a copy number variant (CNV) deletion occurring on the long arm of human chromosome 22, producing a 1.5-3 megabase deletion of 30-50 genes [45–48]. Despite its relative rarity (1 in ∼ 2000-4000 births), the 22qDS conveys the largest known genetic risk factor for developing sporadic schizophrenia and produces cognitive and behavioral phenotypes that have been associated with neurodevelopment disorders such as attention deficit-hyperactivity disorder (ADHD) and autism spectrum disorder [46, 49–51]. Although carriers show high variability in the degree of affected central and peripheral nervous system functions, which are unrelated to deletion size, impairments in attention are commonly observed. Specifically, individuals tend to have problems when attention has to be optimized toward relevant target information and away from irrelevant stimuli [52–55]. Abnormalities in synchronized neural oscillations may contribute to such perceptual and cognitive deficits, as has been hypothesized in schizophrenia, as carriers of the 22qDS have been shown to have a reduced capacity for sensory information to entrain cortical circuits to gamma frequencies [56–58]. These cognitive and neurophysiological features suggest that dysfunctional PVN activity may be a component of the 22qDS pathophysiology, and abnormalities in PVN expression and morphology have been demonstrated in PFC tissue of Df(h22q11/+) mice [59, 60], a model of the 22qDS that also exhibit deficits in focused, but not spatially divided, visual attention [61, 62]. Because the 22qDS conveys such a high penetrance for psychiatric disorders, animal models provide a highly relevant tool for studying the prefrontal mechanisms that may underlie cognitive dysfunction in these conditions.

Although prefrontal PVNs have been shown to support attention, previous studies have primarily evaluated their contributions to detecting and responding to relevant visual cues [10, 11]. Thus far, it has yet to be determined how prefrontal PVNs are involved attentional selectivity (i.e., optimizing focus towards relevant, and identifying and ignoring irrelevant, information), which is highly relevant for their potential contributions to various psychiatric disorders that exhibit deficits in attention. As disrupting PVN activity impairs various PFC-dependent cognitive functions, it is still unclear whether these cells support processing that is consistent across different cognitive tasks, or whether they provide dynamic and flexible support depending on task demands and conditions. Thus, being able to more specifically identify how PVNs contribute to various cognitive functions may yield insight into how their dysfunction may be contributing to the cognitive impairments observed in clinical populations.

In the present study, we assessed mice on a touchscreen-based rodent continuous performance test (rCPT), which was based on human paradigms to provide a translational assessment of cognitive control and focused attention [63, 64]. Importantly, while individuals with conditions such as the 22qDS and schizophrenia are unaffected on tasks that assess simple sustained attention functions such as target detection, paradigms like CPTs include non-target information that must be ignored and reliably produce deficits in the overall ability to discriminate target from non-target stimuli [65–70]. In the rodent version, mice are trained to discriminate between target and non-target images as they are continuously presented individually in the center of a touchscreen. The task progresses through two versions, which retain the same task structure but differ in the probability of target presentations and the number of non-target images, and acute parameters modifications can be made to further tax cognitive load. Our results show that mPFC PVN activity appeared to be adaptive across task training and to acute changes in task demands, consistent with the idea that cortical circuitry including PVNs may be most efficiently deployed in a dynamic, flexible manner [71–75]. We show that poor performance in wild type mice is associated with reduced PVN recruitment and can be rescued by gamma-frequency optogenetic stimulation of PVNs. Finally, we identified abnormal prefrontal PVN activity in the Df(h22q11/+) 22qDS mouse model [49, 76], and were able to alleviate cognitive impairments in these mice using gamma-frequency optogenetic stimulation of PVNs. These data indicate that PVN activity is related to task demands and task performance and may serve as a target for alleviating attention impairments in psychiatric disorders.

## Results

### Prefrontal PVN activity reliably tracks stimulus presentation and correct responses to non-target stimuli

All mice were trained on both versions of rCPT, beginning with the 1 target 1 non-target (1NT) task and followed by the 1 target 4 non-targets (4NT) task, aligning with previous training protocols [63, 77] (Figure 1A). Mice were trained on each task until performance was stable (Hit rate > 0.6; False alarm rate < 0.2; Discrimination sensitivity > 1.5). To assess task-related activity of PVNs we implemented calcium imaging via fiber photometry to record population-level dynamics in the mPFC. Adult male and female PV:Cre mice were unilaterally injected with cre-dependent AAV-syn-FLEX-jGCaMP7f-WPRE (GCaMP7f, [78] into the mPFC, and implanted with a fiber optic probe for light delivery and collection prior to task training (Figure 1A). PVN signals were assessed during each behavioral response (Hit, Miss, False Alarm (FA), and Correct Rejection (CR)) on the final training session for both the 1NT and 4NT tasks (Figure 1B). We first looked at the change in PVN activity at stimulus presentation, and found no differences in PVN recruitment when mice would go on to make either response type on the 1NT (Figure 1C), or the 4NT (Figure 1D), task. Next, we aligned PVN signals to the time a response was recorded (i.e., a physical screen touch or removal of stimulus for non-response measures) (Figure 1E, also see Supplementary Figure 1A-B). We compared responses to both target and non-target images and found significantly elevated PVN activity when animals made a correct rejection to non-target images during both the 1NT (Figure 1F) and 4NT task (Figure 1G). We next asked if there were relationships between PVN signals and specific task events, such as image type or response outcome (i.e., correct or incorrect). On the 1NT task, we observed a positive correlation between PVN signals at stimulus presentation when animals would go on to make a correct response to either a target or non-target (Figure 1H). Furthermore, relationships were observed for both response outcomes on 4NT task, where PVN signals prior to making correct responses were positively correlated, as were signals prior to making incorrect responses (Figure 1I). Interestingly, there were no relationships between PVN signals at the time when a response was recorded on either task (Figure 1J-K).

**Figure 1.**
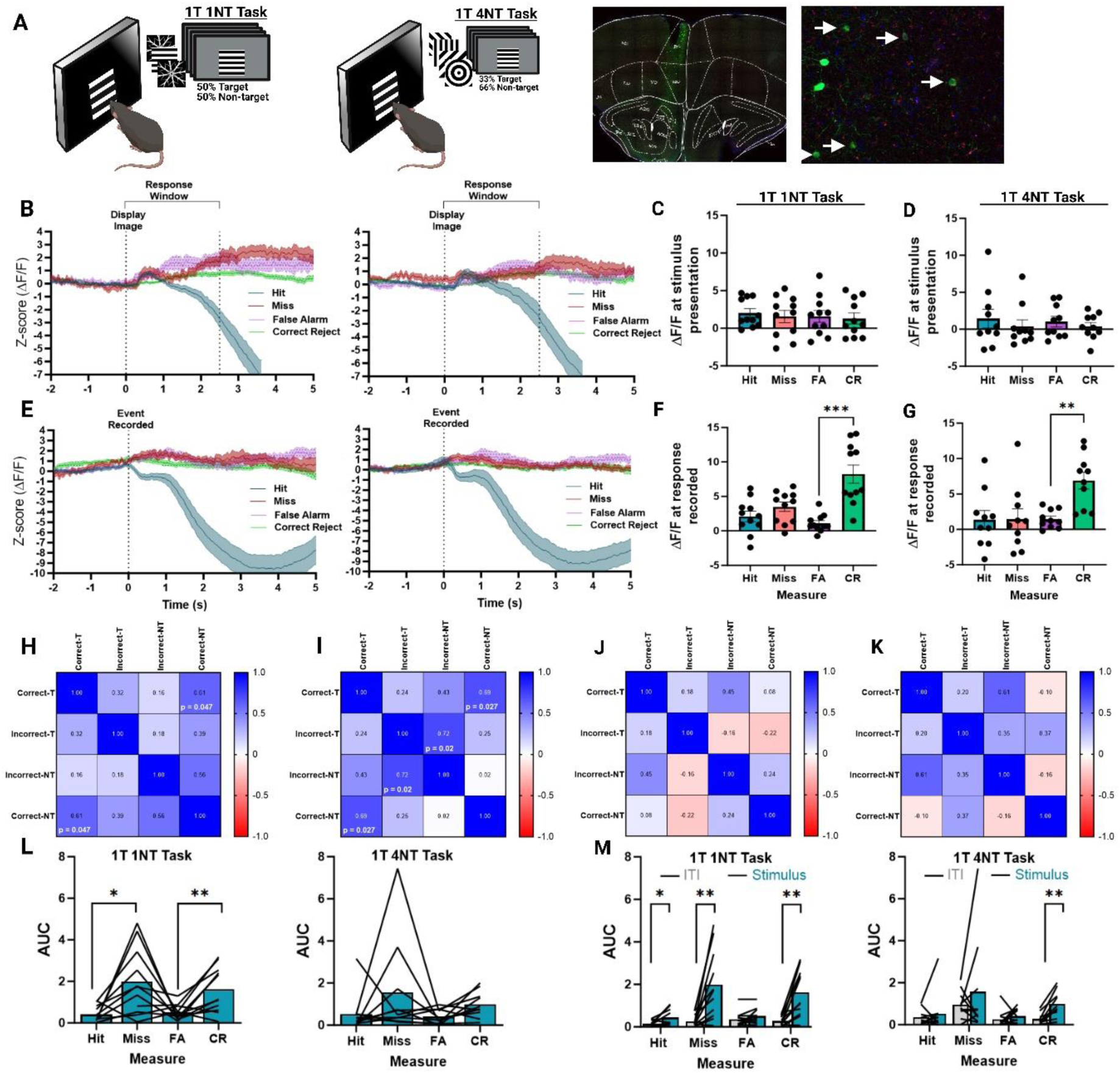
Prefrontal PVN activity reliably tracks stimulus presentation and correct responses to non-target stimuli. **A.** The 1 Target, 1 Non-target (1NT) task (left) and 1 Target, 4 Non-target (4NT) task (Left). A representative image showing GCAMp7f expression and fiber tract in the mPFC (left), alongside a higher power image showing examples of co-localization of GFP-tagged GCaMP7 (green) with PV (red). **B.** Averaged change in fluorescence (ΔF/F) from prefrontal PVN signals time-locked to stimulus presentation on the 1NT task (left) (n = 11 mice; male (n = 6), female (n = 5)) and 4NT task (right). There were no significant effects of sex or interactions (p > 0.05), so data were pooled across sexes (n = 10 mice; male (n = 5), female (n = 5)). **C.** The average ΔF/F from PVN signals calculated from baseline intertrial interval (ITI) activity immediately after stimulus presentation on the 1NT task. **D.** The average ΔF/F immediately after stimulus presentation on the 4NT task. **E.** Averaged ΔF/F from prefrontal PVN signals time-locked to moment a response was recorded on the 1NT task (left) and 4NT task (right). **F.** The average ΔF/F immediately prior to stimulus response on the 1NT task. There was a significant difference in ΔF/F across behavioral measures (One-way repeated measures (RM) ANOVA: F_3, 30_ = 14. 208, p < 0.001), specifically during Correct Rejections compared to False Alarms (p < 0.001). **G.** The average ΔF/F immediately prior to stimulus response on the 1NT task. There was a significant difference in ΔF/F across behavioral measures (One-way RM ANOVA: F_3, 27_ = 6.702, p = 0.002), specifically during Correct Rejections compared to False Alarms (p = 0.003). **H-I.** A correlation matrix analyzing the ΔF/F from PVN signals during initial stimulus presentation preceding correct and incorrect responses to target and non-target stimuli on the 1NT (**H**) and 4NT task (**I**). R^2^ values are shown in black text and significant p values are displayed in white text where appropriate. **J-K.** A correlation matrix analyzing the ΔF/F from PVN signals at the time correct and incorrect responses to target and non-target stimuli were recorded on the 1NT (**J)** and 4NT (**K)** task. R^2^ values are shown in black text. **L.** The area under the curve (AUC) for PVN signals during each response on the 1NT (left) and 4NT task (right). There was a significant effect of response type on the 1NT task (One-way RM ANOVA: F_1.546, 15.46_ = 5.92, p = 0.017), with greater PVN activity during Misses compared to Hits (p = 0.013), and Correct Rejections compared to False Alarms (p = 0.006). There were no significant effects of response type on the 1NT task (F_1.355, 12.195_ = 1.736, p = 0.217). **M.** The area under the curve (AUC) for PVN signals comparing the ITI and stimulus presentation periods on the 1NT (left) and 4NT task (right). There was a significant effect interaction between task period and response type on the 1NT task (Two-way RM ANOVA: F_1.572, 15.719_ = 6.330, p = 0.014), with significant PVN recruitment during Hit (p = 0.022) Miss (p = 0.004), and Correct Rejections (p = 002). There was no significant effects of response type on the 1NT task (F_1.355, 12.195_ = 1.736, p = 0.217). On the 4NT task, there was a significant effect interaction between task period and response type (Two-way RM ANOVA: F_1.572, 15.719_ = 6.330, p = 0.014), where significant PVN recruitment was observed during Correct Rejections only (p = 0.001). Error bars indicate mean ± SEM. *** p < 0.001, ** p < 0.01, * p < 0.05.

To further quantify PVN activity, we compared the area under the curve (AUC) of PVN signals prior to each response. While PVN activity was significantly higher for non-response outcomes (i.e., a Miss or Correct rejection) during target and non-target presentations on the 1NT task, there were no differences across measures on the 4NT task (Figure 1L). Lastly, we evaluated the AUC of PVN signals during stimulus presentation compared to baseline intertrial interval (ITI) activity across behavioral measures. PVN actually was significantly elevated during most response outcomes on the 1NT task, while increases in PVN activity on the 4NT were isolated to correct rejections (Figure 1M). Similarly, an analysis of the maximum ΔF/F during stimulus presentation revealed somewhat different profiles across the two tasks, where activity was generally increased across measures on the 1NT task but specific to non-response outcomes on the 4NT task (Supplementary Figure 1 C-D). Additionally, a mixed models linear cross-task analysis revealed a significant difference in max PVN ΔF/F during stimulus presentation, suggesting that PVN recruitment may become refined with extensive task training (F_1, 76_ = 5.686, p = 0.0196) (Supplementary Figure 1E). Lastly, a consistent finding across the task versions was a significant suppression of PVN activity following a hit response, which was the only response that resulted in reward delivery, consistent with previous studies describing reduced prefrontal PVN activity at reward delivery [79–82]. Moreover, PVN activity was significantly higher following an incorrect response to non-target stimuli (false alarm) compared to a correct rejection, suggesting a bi-directional modulation of prefrontal PVN signalling following reward and “punishment” (i.e., absence of reward) (Supplementary Figure 1E).

### Prefrontal PVN activity is altered by acute task demands and associated with task performance

Although we observed some subtle differences in PVN signal patterns between the two versions of the CPT, the inclusion of the additional non-target images and the reduced target probability did not seem to be a large enough deviation from task structure and demands to induce distinct activity profiles. Therefore, we next taxed stimulus detection and discrimination by reducing the stimulus presentation period across testing sessions (2.5s to 0.5 and 0.2s) and assessed whether acute changes in task demands were represented in PVN activity (Figure 2A-B, Supplementary Figure 2). To assess pre-stimulus activity, we looked at the maximum ΔF/F immediately prior to stimulus presentation. There was a significant shift in the PVN activity profile during the ITI period, which was characterized by a significant increase in activity preceding a screen response (i.e., hit or false alarm), as well as a decrease in activity preceding a miss, particularly at the lowest stimulus duration (Figure 2C). Further, there was a significant reduction in the ΔF/F of PVN signals at stimulus presentation only when animals made a Correct Rejection (Figure 2D). Interestingly, there was also a change in PVN signals following correct responses, where activity increased during the post stimulus period following a Hit and decreased following Correct Rejections (Figure 2E). Together, these data suggest that acute changes in task demands can modify the recruitment of PVN activity, here potentially reflecting alterations in vigilance requirements and post-response updating.

**Figure 2.**
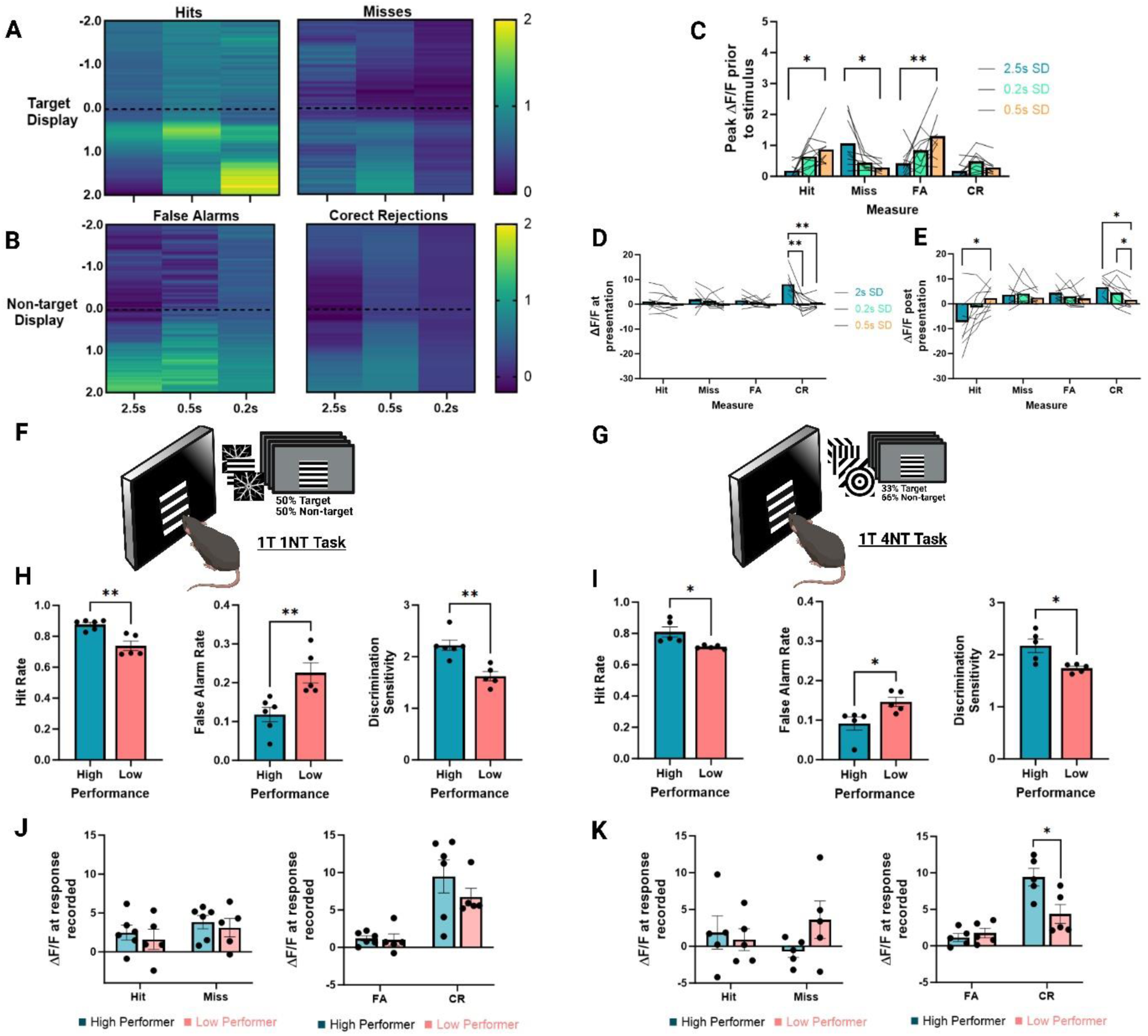
Prefrontal PVN activity is altered by acute task demands and associated with task performance. **A.** Averaged ΔF/F from prefrontal PVN signals during correct (left) and incorrect (right) responses to target stimuli on each stimulus duration (n = 9 mice). **B.** Averaged ΔF/F from prefrontal PVN signals during correct (left) and incorrect (right) responses to non-target stimuli on each stimulus duration (n = 9 mice). **C.** The maximum ΔF/F peak time-locked to the ITI period preceding stimulus presentation for each measure across three stimulus duration lengths (2.5s, 0.5s, 0.2s). There was a significant interaction between stimulus duration and PVN signal across measures (F_2.492, 19.934_ = 7.269, p = 0.001). *Post hoc* tests revealed a significant increase in PVN activity during the lowest stimulus duration preceding a Hit (p = 0.046) and False Alarm (p = 0.003). Additionally, there was a significant decrease in PVN activity preceding a Miss (p = 0.011). **D.** The average ΔF/F from prefrontal PVNs during stimulus presentation for each behavioral measure across stimulus durations. There was a significant interaction between behavioral measure and stimulus duration (Two-way ANOVA: F_2.108, 16.86_ = 6.249, p = 0.0086), and *post hoc* tests showed a specific decrease in PVN activity during Correct Rejections at 0.5s (p = 0.008) and 0.2s (p = 0.005) durations. **E.** Average ΔF/F from prefrontal PVNs during stimulus presentation for each behavioral measure across stimulus durations. There was a significant interaction between behavioral measure and stimulus duration (Two-way ANOVA: F_2.551, 20.41_ = 6.583, p = 0.004), and *post hoc* tests showed a specific increase in PVN activity following Hit responses at 0.2s (p = 0.047), and a significant decrease in activity following Correct Rejections at 0.5s (p = 0.045) and 0.2s (p = 0.015) durations. **F-G.** The 1NT (left) and 4NT (right) task. Figures shown below the corresponding diagram presents data collected using the corresponding task. **H.** Separation of mice into high and low performance groups for the three primary behavioral measures on the 1NT task. Low performing mice were significantly worst across measures compared to high performers (One-way ANOVA: F_1, 9_ = 17.282, p = 0.002; Hit Rate (HR), p = 0.002; False Alarm Rate (FAR), p = 0.008; Discrimination sensitivity (d’), p = 0.002). **I.** Separation of mice into high and low performance groups for the three primary behavioral measures on the 4NT task. Low performing mice were significantly worst across measures compared to high performers (One-way ANOVA: F_1, 8_ = 13.286, p = 0.007; Hit Rate (HR), p = 0.018; False Alarm Rate (FAR), p = 0.028; Discrimination sensitivity (d’), p = 0.013). **J.** Average ΔF/F from prefrontal PVNs in high and low performers at the time of response to target (left) and non-target (right) stimuli on the 1NT task. **K.** Average ΔF/F from prefrontal PVNs in high and low performers at the time of response to target (left) and non-target (right) stimuli on the 4NT task. There was a significant interaction between ΔF/F and performance for responses to non-targets (F_1, 8_ = 8.438, p = 0.020), specifically between high and low performers when making Correct Rejections (p = 0.011). Error bars indicate mean ± SEM. ** p < 0.01, * p < 0.05.

Lastly, we asked whether PVN activity was associated with proficiency on both CPT tasks. To do so, animals were classified as high or low performers using a median split analysis based on their response rates to either target or non-target stimuli on both paradigms (Figure 2F-G). Across performance measures, animals designated as low performers were significantly worse than their high-performance counterparts on both the 1NT (Figure 2H) and 4NT task (Figure 2I). While there was no interaction between task performance and PVN activity during the 1NT task (Figure 2J), animals that exhibited a better false alarm rate on the 4NT task showed significantly higher activity during correct rejections than lower performing animals (Figure 2K).

### Frequency-specific activation of prefrontal PVNs differentially modulates local oscillatory activity

To examine the causal contributions of prefrontal PVNs to task performance, we next used *in vivo* optogenetics to modulate PVN activity across the two task variants. Mice were generated by crossing PV:Cre mice with lines ubiquitously expressing either GFP-tagged inhibitory archaerhodopsin (Arch) or excitatory channelrhodopsin-2 (ChR2) to generate offspring with selective PVN-opsin expression (PV:Arch and PV:ChR2 mice, respectively) [83]. We first aimed to confirm that frequency specific activations of PVNs (i.e., 5hz and 30hz) could modulate the local oscillatory profile *in vivo*. Previous studies have reported bi-directional effects of PVN stimulation on cognition, where low gamma (30-40hz) stimulation has been shown to improve task performance while lower theta stimulation (5hz) impaired performance [6, 10]. Local field potentials (LFPs) were collected in anesthetized mice in response to blue-light excitation of local PVNs in the mPFC (Figure 3A). Stimulation of PVNs at 5hz was characterized by significant increases in evoked power (EP) across all frequency bins (Figure 3B), as well as a significant increase in intertrial coherence (ITC) at the low- and high-gamma frequency bins (Figure 3C). In contrast, 30hz stimulation of PVNs within the mPFC resulted in a gamma-specific enhancement of EP and ITC, which was observed solely within the low-gamma frequency range (Figure 3B-C). To further quantify the overall oscillatory profile within the optogenetic stimulation period (i.e., 0 to 2s), the scaled power was calculated across the oscillatory spectrum for each of the stimulation conditions (5hz, 30hz and no light). As expected, 30hz stimulation of PVNs enhanced the local gamma power specifically within the low-gamma band (i.e., 26 – 34hz) (Figure 3E). Although 5hz stimulation of PVNs resulted in no significant alterations in the scaled power of oscillations beyond the low frequencies, oscillatory peaks at 5hz intervals were clearly observed (see red line in Figure 2D). Thus, in comparing the two optogenetic stimulation conditions on the overall oscillatory profile, the 30hz stimulation specifically increased mPFC gamma frequency band activity whereas stimulating PVNs at 5hz altered the local oscillatory profile across a range of frequencies.

**Figure 3.**
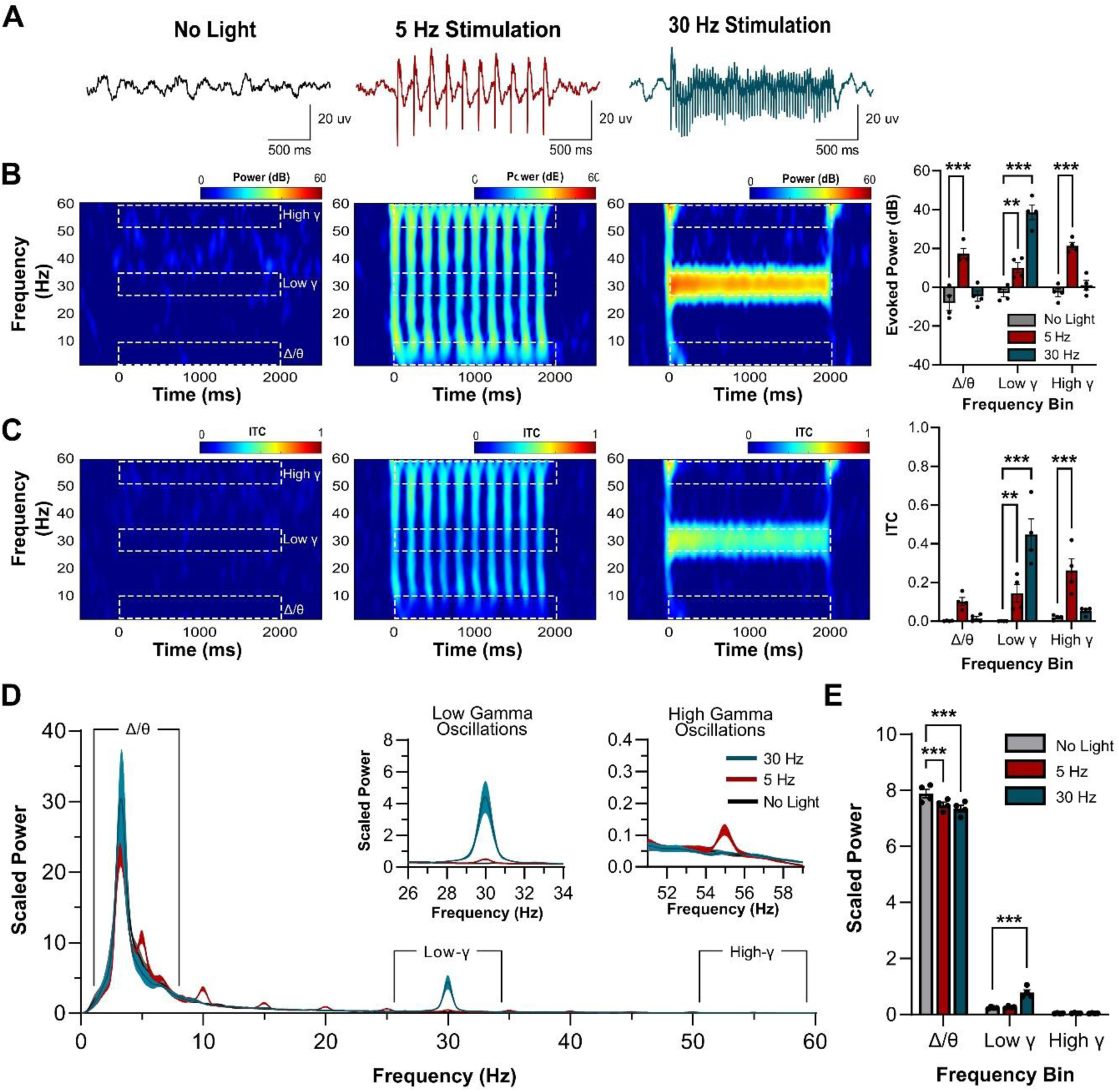
Frequency-specific activation of prefrontal PVNs differentially modulates local oscillatory activity. **A.** Average local field potentials in response to no light (black), 5hz (red) and 30hz (blue) optogenetic stimulation within the mPFC (n = 4 mice). **B.** Heat maps plot the group average of phase-locked evoked power (EP) in response to frequency-specific activation (no light = left panel; 5hz stimulation = middle panel; 30hz = right panel). Evoked power was quantified across three frequency ranges (i.e., delta/theta (Δ/Ѳ): 1 – 9hz; Low-γ: 26 – 34hz; High-γ: 51 – 59hz) during the presentation of the optogenetic stimuli (0 and 2000 ms; denoted by the white boxes on all heat maps). A two-way RM ANOVA revealed a significant interaction between frequency range (i.e., Δ/Ѳ, Low-γ and High-γ) and optogenetic activation frequency (i.e., no light, 5hz and 30hz) (F_4, 12_ = 34.30, p < 0.001). Post-hoc analyses revealed a significant increase in evoked power across all frequency ranges (Δ/Ѳ: p < 0.001; Low-γ: p = 0.007; High-γ: t_4_ = 6.683, p < 0.001) following the activation of PVNs at 5hz, while only Low-γ showed a significant increase in EP following 30hz optogenetic activation (Low-γ: p < 0.001). **C.** Phase-locked intertrial coherence (ITC) is plotted in the heat maps in response to frequency-specific activation (no light = left panel, 5hz stimulation = middle panel; 30hz = right panel). Consistent with the analysis performed on EP, ITC was quantified across 3 frequency ranges (i.e., Δ/Ѳ, Low-γ and High-γ) during optogenetic stimulation. A two-way RM ANOVA revealed a significant interaction between frequency range and activation frequency (F_4,12_ = 26.70, p < 0.001). Post-hoc analyses revealed a significant increase in ITC within both the gamma frequency ranges (Low-γ: p = 0.009; High-γ: p < 0.001) following the 5hz stimulation. Activation of PVNs at 30hz resulted only in a significant increase in ITC within the Low-γ frequency range (p < 0.001). **D.** Average scaled power during frequency specific activation (i.e., 0 to 2000 ms) recorded from the mPFC during 5hz (red) and 30hz (blue) activation as well as no light (black) (shade indicates SEM). Inserts show the scaled power oscillatory profile in the Low-gamma (Low-γ) and High-gamma (High-γ) frequency ranges. Bar graphs on the right show average scaled power within each frequency bin. A two-way RM ANOVA revealed a significant interaction of frequency bin by activation frequency (F_4, 12_ = 29.79, p < 0.001). In the low frequency bin (i.e., Δ/Ѳ), there was a significant decrease in scaled power following the 5hz activation (p < 0.001) and the 30hz activation (p < 0.001) when compared to the no light condition. Within the Low-gamma frequency range there was a significant increase in scaled power following 30hz activation (p < 0.001). There was no change in scaled power within the High-gamma frequency range following either activation frequency (p > 0.05). Error bars indicate mean ± SEM. *** p < 0.001, ** p < 0.01, * p < 0.05.

### Optogenetic disruption of prefrontal PVNs impairs target responding during the 1NT Task

To assess the effects of PVN disruption on the 1NT task, we implanted male and female PV:Arch mice with bilateral fiber optic probes to deliver light to the mPFC. Once mice were trained to the criterion outlined above, light (550nm, constant, ̴ 2.5mW/side) was delivered pseudorandomly on half of image presentations during the testing session (Figure 4A-B). Inhibition of PVNs significantly reduced overall discrimination (Discrimination sensitivity) in both male and female mice, which was driven by a reduction in target responding (Hit rate) (Figure 4C). False alarm rate, response bias (Figure 4C), and latency measures (Supplementary Figure 3A-B) were unaffected, and there were no effects of sex on any measures. Additionally, we found that PVN inhibition during the ITI or stimulus presentation, individually, had no effect on performance (Supplementary Figure 4). Although a number of studies have reported no off-target behavioral effects of light delivery to the mPFC (e.g., non-opsin mediated effects), the lack of behavioral effects observed here also suggest the presence of the light itself is not altering performance. To activate PVNs at either theta (5hz) or gamma (30hz) frequencies, male and female PV:ChR2 mice underwent the identical surgical and behavioral protocols. After performance was stable, mice received counterbalanced sessions with optogenetic stimulation (465nm, 5ms, ̴ 1mW/side) at either 5hz or 30hz. As with Arch, 5hz activation significantly reduced target responding in both male and female mice (Figure 4D), with no effects of stimulation on task latency measures (Supplementary Figure 3C-D). Thus, perturbing prefrontal PVN activity induced a consistent deficit in responding to the target stimulus on the 1NT task. Interestingly, 30hz stimulation induced a more conservative style of responding in female mice only, characterized by a reduction in the tendency to respond to the screen (Response bias, Figure 4E). Response and reward collection latencies were unaffected (Supplementary Figure 3E), and there were no sex differences in baseline performance between sexes.

**Figure 4.**
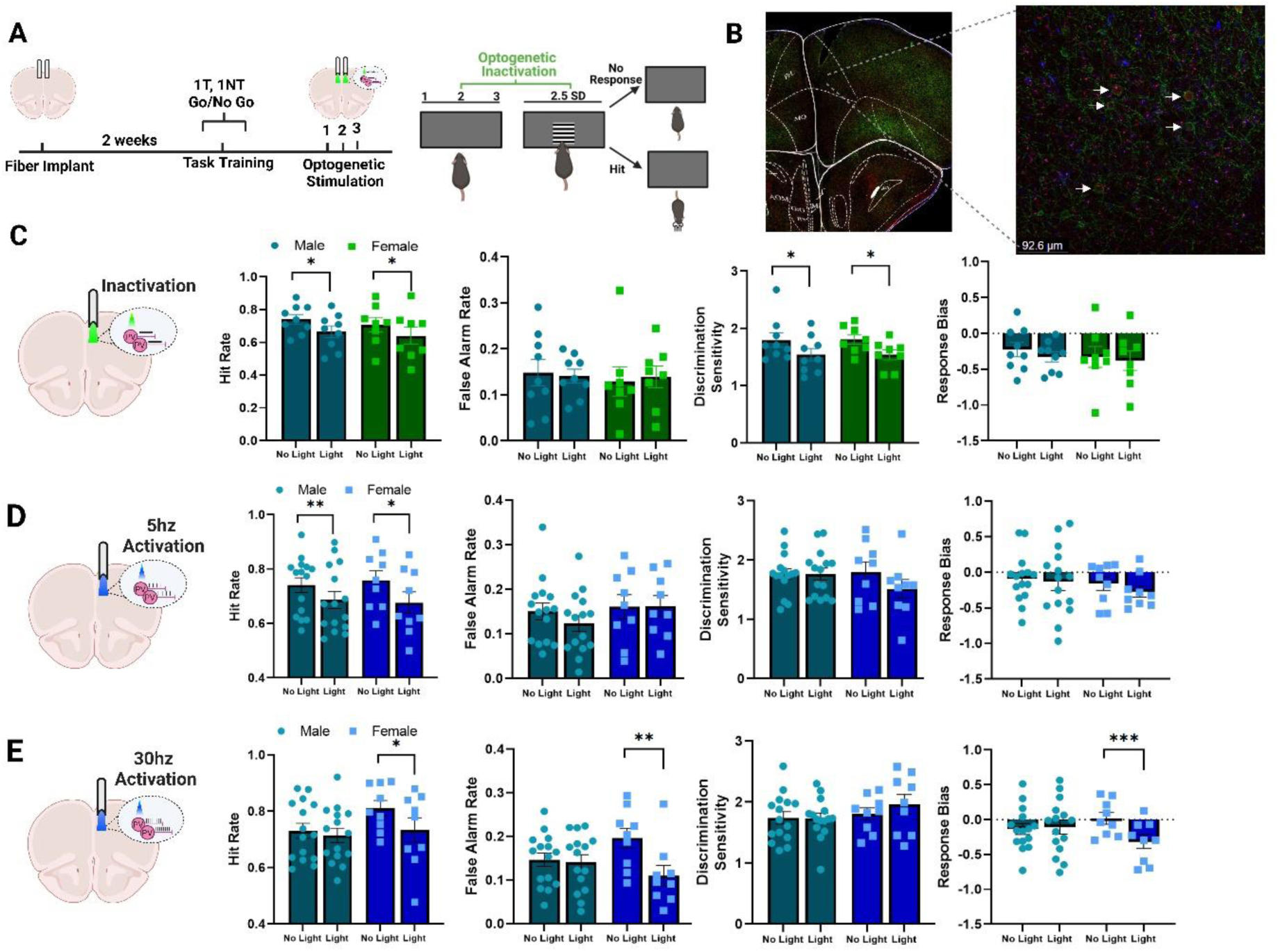
Optogenetic disruption of prefrontal PVNs impairs target responding during the 1NT task. A. Experimental timeline for optogenetic manipulations on the 1NT task. B. Fiber optic tracts and expression of GFP (green) with PV (red) in a coronal section of mPFC (left), alongside a high-power image (right, white arrows indicate examples of co-expression, 40x). C. Optogenetic inactivation of prefrontal PVNs significantly reduced the hit rate (Two-way RM ANOVA: F_1, 15_ = 17.742, p < 0.001) and discrimination sensitivity (F_1, 15_ = 16.819, p < 0.001) in both male (p = 0.013; p = 0.024, respectively) and female (p = 0.027; p = 0.018) mice. There were no significant effects of optogenetic inhibition on false alarm rate (F_1, 15_ = 0.011, p = 0.920) or response bias (F_1, 15_ = 3.935, p = 0.066), nor effects of sex or interactions for any measure. n = 9 male, 8 female mice. D. 5hz activation of PVNs significantly reduced the hit rate (Two-way RM ANOVA: F_1, 22_ = 19.130, p < 0.001) in both male (p = 0.007) and female (p = 0.021) mice. There were no significant effects of optogenetic inhibition on false alarm rate (p = 0.238), discrimination sensitivity (p = 0.059) or response bias (p = 0.323), nor effects of sex or interactions for any measure (n = 15 male, 9 female mice). E. 30hz activation of PVNs did not significantly affect any measures in male mice. However, overall ANOVAs showed significant reductions in hit rate (Two-way RM ANOVA: F_1,22_ = 5.442, p = 0.029), false alarm rate (F_1,22_ = 12.427, p = 0.002; opto x sex interaction, p = 0.005), and response bias (F_1, 22_ = 11.002, p = 0.003; opto x sex interaction p < 0.001), illustrating a general trend in reduced responding in female mice (p = 0.035; p = 0.002; p < 0.001, respectively). There were no main effects of sex for any measure. Error bars indicate mean ± SEM. *** p < 0.001, ** p < 0.01, * p < 0.05.

### Gamma stimulation of prefrontal PVNs improves attention in low performing mice on the 4NT task

After observing that PVN perturbation induced performance deficits on the 1NT task, we next investigated the effects of manipulating PVN activity on the 4NT task. PV:ChR2 mice were trained on the 4NT rCPT until performance was stable and optogenetic stimulation was delivered in an identical manner to that for the 1NT task (Figure 5A). Interestingly, PVN perturbation with 5hz stimulation induced deficits in discrimination sensitivity and non-target identification on this version of the CPT, while target responding was unaffected (Figure 5B). Furthermore, 30hz stimulation significantly enhanced overall discrimination and non-target identification (Figure 5C), and neither manipulation impacted any latency measures (Supplementary Figure 5). Importantly, the observation of bi-directional effects with the same light wavelength suggests a lack of non-specific improvements or impairments in behavior as a result of light delivery. Lastly, a mixed models linear cross-task analysis revealed a significant interaction for discrimination sensitivity (task version x optogenetic stimulation x frequency; F_1,11_ = 5.414, p = 0.0369), suggesting a sensitivity of PVN-contributions to distinct task requirements. Although the PVN GCaMP signals appeared to be relatively similar across the two task versions, the distinct effects of the gamma stimulation suggest a functionally different contribution of PVNs to performance across task versions.

**Figure 5.**
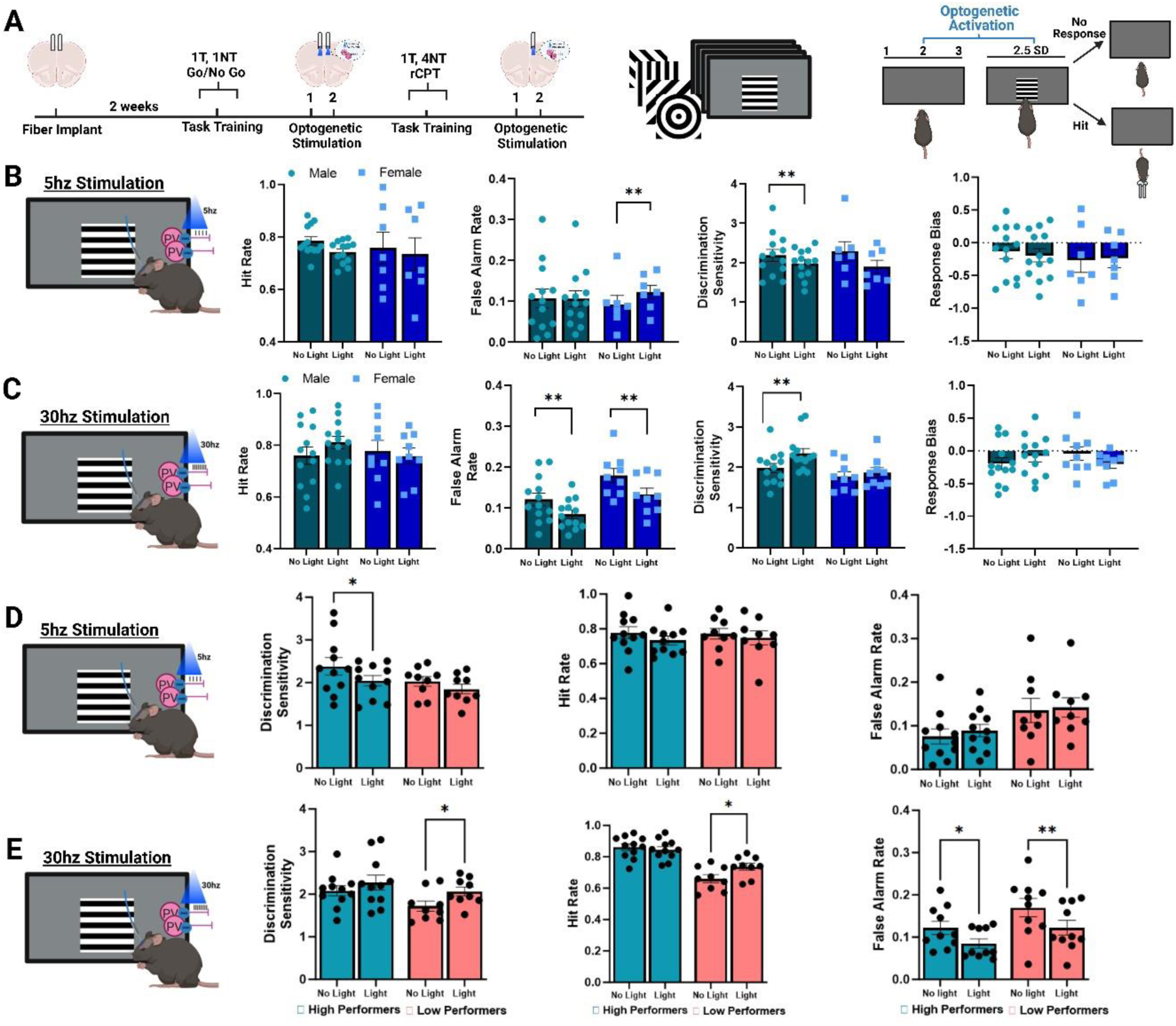
Gamma stimulation of prefrontal PVNs improves attention in low performing mice on the 4NT task. A. Experimental timeline for optogenetic activation of PVNs during the 4NT rCPT. Two male mice and did not complete 4NT training due to headcap damage. Additionally, two more female mice were unable to complete the 5hz stimulation session due to headcap damage. (n = 13 male mice, 9 female mice). B. 5hz activation of PVNs significantly increased false alarm rate (F_1, 18_ = 4.635, p =0.045; opto x sex interaction p = 0.041) in female (p = 0.045) but not male (p = 0.966) mice. Additionally, 5hz activation decreased discrimination sensitivity (F_1, 18_ = 12.174, p = 0.002), though this was selective to male mice (p = 0.031) and not females (p = 0.061). There were no significant effects of 5hz stimulation or sex on hit rate (F_1, 18_ = 3.920, p = 0.063) or response bias (F_1, 18_ = 0.172, p = 0.683). C. 30hz activation of PVNs significantly reduced false alarm rate (F_1, 20_ = 25.578, p < 0.001; effect of sex p = 0.010, no interaction) in both male (p = 0.004) and female (p = 0.007) mice. Additionally, 30hz activation increased discrimination sensitivity (F_1, 20_ = 8.510, p = 0.009, effect of sex p = 0.040, no interaction) specifically in male mice (p = 0.003). There were no significant effects of 30hz stimulation or sex on any additional measures. D. PV:ChR2 mice (n = 20) were classified as high (blue) or low (red) performers using a median split analysis. There was a significant interaction of stimulation x performance measure (F _(1.65, 29.64)_ = 300.17, p > 0.001), where *post hoc* tests showed a significant reduction in discrimination sensitivity in high (p = 0.014), but not low performers (p = 0.138). E. The effect of modulating PVN activity with 30hz stimulation was assessed as a function of performance level. There was a significant opto x performance x behavioral measure interaction (F_2, 32_ = 13.689, p > 0.001). 30hz activation significantly increased discrimination sensitivity, in low (p = 0.017) but not high performers (p = 0.610), and hit rate in low (p = 0.017) but not high performers (p = 0.610). 30hz activation significantly improved false alarm rate in both high (p = 0.015) low-performing mice (p = 0.004). Error bars indicate mean ± SEM. ** p < 0.01, * p < 0.05.

After observing that greater PVN recruitment was associated with better non-target identification on the 4NT rCPT (Figure 1K), we wanted to assess whether baseline rCPT performance affected the efficacy of optogenetic stimulation. Moreover, various studies testing attention-enhancing pharmacological compounds in rodents show that improved performance is often restricted to animals who perform the given task more poorly [84–87]. If less optimized PVN activity is present in lower performing animals and can be alleviated with artificial gamma stimulation, perhaps a similar approach could be taken for targeting abnormalities in PVNs present in psychiatric disorders. To investigate this question, we categorized the mice as high or low performers using a median split analysis based on their averaged performance for each measure across the 30hz and 5hz “No Light” conditions. We observed that while 5hz stimulation impaired discrimination sensitivity only in high-performing mice, 30hz stimulation improved discrimination sensitivity only in low-performing mice (Figure 5D-E, respectively). Interestingly, improved discrimination was the result of optimizing both the hit rate and false alarm rate, illustrating that hit rate could only be improved via 30hz stimulation in poorer performing animals (Figure 5E). These data, in combination with the interaction observed between higher PVN activity and better task performance, indicate that stimulating PVNs at the gamma frequency has the potential to improve cognition in poorly performing mice.

### Learning amplifies PVN activity and is absent in a mouse model of the 22q11.2 microdeletion syndrome that exhibits attention deficits

The Df(h22q11/+) model (22q) provides a well characterized tool to assess the involvement of PVNs in abnormal attention, as these mice exhibit rCPT deficits [62] and reduced PVN-marker expression in the mPFC [59, 60]. We first replicated the attention deficit in male mutant mice and confirmed female mutants were also impaired on the 1NT rCPT [62]. 22q mice displayed a significant reduction in discrimination sensitivity across task training, and an associated reduction in target detection (Supplementary Figure 6A-B). Of note, reduced discrimination sensitivity reflects the performance deficit that is observed in human patients with 22qDS and psychiatric disorders on CPT paradigms [65–70, 88].

To assess the real-time function of prefrontal PVNs during attention, which to our knowledge has not been assessed in any models of the 22qDS, we crossed hemizygous Df(h22q11.2/+) mice with homozygous PV:Cre mice to generate wild-type and 22q:PV-Cre offspring. 22q and wild-type mice were unilaterally injected with the cre-dependent GCaMP7f calcium biosensor [78] into the mPFC and implanted with a fiber optic probe for light delivery and collection. On the final 1NT task training session, 22q mice exhibited significantly lower prefrontal PVN activity during the ITI prior to and during the stimulus presentation when making a Hit response (Figure 6A). There were no differences between groups during Miss or False Alarm responses (Figure 6B-C), but 22q mice exhibited significantly lower PVN activity during the correct rejections, together showing reduced PVN recruitment in 22q mice during correct responses across image types (Figure 6D). We next sought to evaluate whether prefrontal PVN activity was modulated across task acquisition, and whether abnormalities in prefrontal PVN organization may contribute to the observed behavioral deficit in 22q mice. Because blunted PVN activity was observed during correct responses in 22q mice, we analyzed signals across learning for correct and incorrect responses on the first session and the last session when animals reached criteria. In wild-type mice, PVN activity was significantly higher on the final training session during correct, but not incorrect, responses (Figure 6E-H). Opposingly, 22q mice did not show a modification of PVN signal across learning for any behavioral measure. Thus, we demonstrate that the activity of prefrontal PVNs is modulated across learning, represented in stronger signal recruitment during correct responses. Although this pattern was absent in 22q mice, the functional capacity of prefrontal PVNs in the context of psychiatric disorders is an important question, as reports of lower PV expression and/or numbers of PV cells in the PFC questions the viability of targeting this population to alleviate cognitive impairments [19, 25, 28, 34, 35, 89]. While PVN activity is heavily blunted in the 22q mice, we observed remnants of functional dynamics, including the characteristic suppression of PVN activity following a rewarded Hit (Supplementary Figure 6C-D). Thus, we did not observe a complete absence of activity dynamics, suggesting that this population retains some functional characteristics and may be a target for direct intervention.

**Figure 6.**
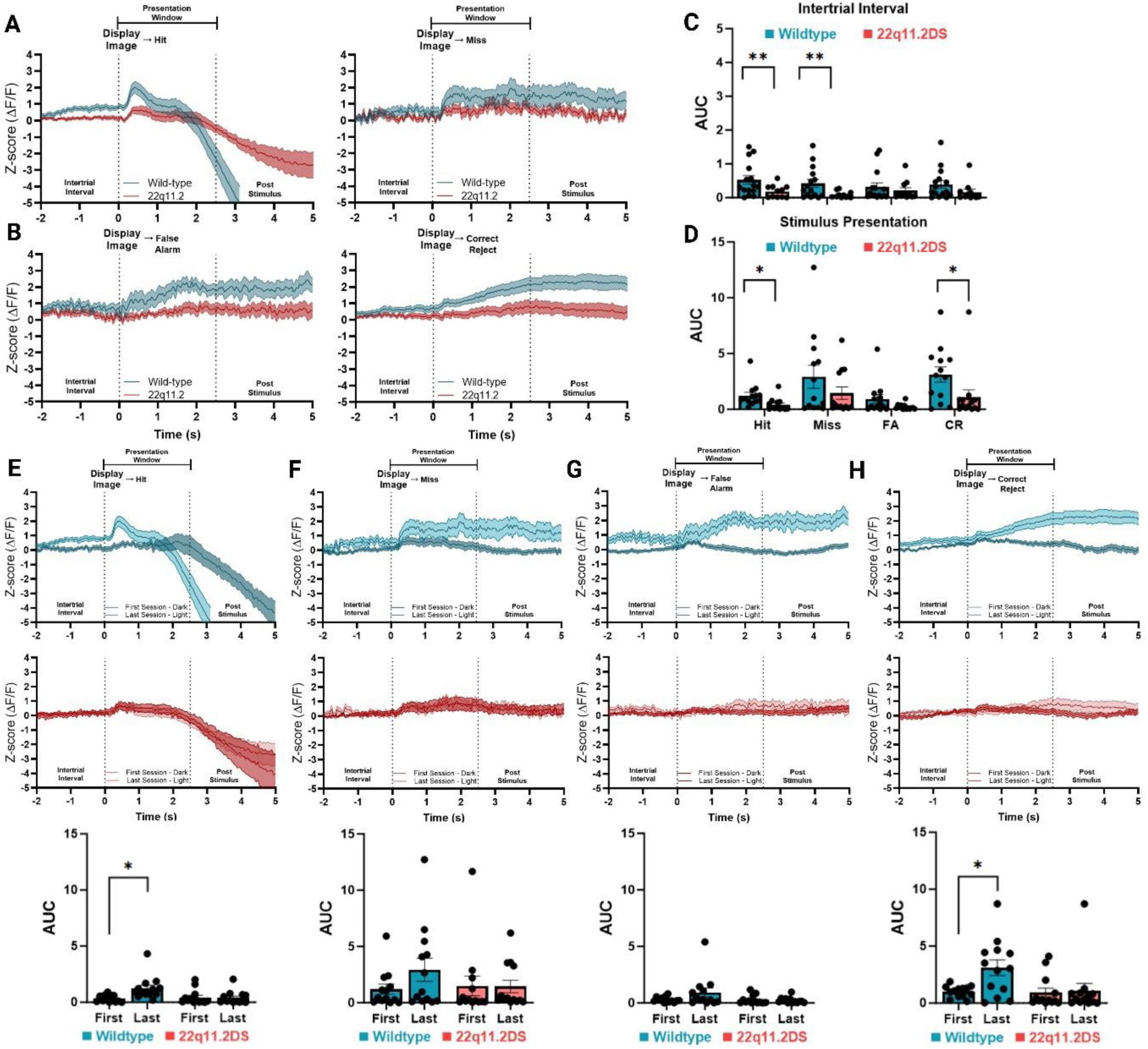
Learning amplifies PVN activity and is absent in a mouse model of the 22q11.2 microdeletion syndrome that exhibits attention deficits. **A.** Averaged ΔF/F from prefrontal PVN signals during responses to target stimuli on during the last session of 1NT training. n = 13 wild-types (WT; blue: 9 males, 4 females) and 13 22q11.2 (22q; red: 7 males, 6 females) mice. There were no significant effects or interactions of sex (p > 0.05), so data were pooled across sexes. **B.** Averaged ΔF/F from prefrontal PVN signals during responses to target stimuli on during the last session of 1NT training. **C.** AUC analysis comparing prefrontal PVN activity between WT and 22q mice during the ITI period preceding stimulus presentation. 22q mice display significantly lower PVN activity preceding target presentations (Two-way RM ANOVA: F_1, 22_ = 10.977, p = 0.003) both when an animal would go on to make a Hit (p = 0.009) or Miss (p = 0.020). **D.** AUC analysis comparing prefrontal PVN activity between WT and 22q mice during the stimulus presentation leading to a response. 22q mice display significantly lower PVN activity during correct responses (Two-way RM ANOVA: F_1, 22_ = 5.213, p = 0.032) both when an animal would go on to make a Hit (p = 0.016) or Correct Rejection (p = 0.043). **E-H.** Comparison of the average ΔF/F from prefrontal PVN during the first training session (Light colors) compared to the final training session where criterion was reached (Dark colors). **Top Row:** Prefrontal PVN activity in WT (blue) mice. From left to right figures: Hit, Miss, False Alarm, Correct Rejection. **Middle:** Prefrontal PVN activity in WT (blue) mice. From left to right figures: Hit, Miss, False Alarm, Correct Rejection**. Bottom:** AUC analysis for PVN activity on the first and last session of training. There was a significant interaction between session and genotype for correct (Two-way ANOVA: F_1, 24_ = 7.302, p = 0.012) but not incorrect responses (Two-way ANOVA: F_1, 24_ = 0.963, p = 0.336). *Post hoc* tests show a significant increase in PVN activity during Hits (p = 0.012) and Correct Rejections (p = 0.011) in WT but not 22q mice.

### Gamma stimulation of PVNs improves non-target responding and rescues deficits in discrimination sensitivity in a mouse model of 22q11.2 microdeletion syndrome

Previous studies have shown that directly targeting prefrontal PVNs is effective for restoring aspects of cognition in rodent models of psychiatric disorders, potentially through the restoration of gamma activity [6, 9, 11, 90]. To target PVNs in the 22qDS model, we injected AAV-CAG-DIO-ChR2(H134R) bilaterally into the mPFC of male and female 22q:PV-Cre mice and wild-type PV:Cre controls (Figure 7A-B). Mice were assessed over 3 sessions on the 1NT task, where we again observed attention deficits in 22q mice (Figure 7C-D), with no differences across genotype or sex in latencies to respond or collect reward (Supplementary Figure 7). On the 4th session, 30hz optogenetic stimulation was applied pseudorandomly throughout the session using the same protocol as previously described. Here, discrimination sensitivity was significantly lower for 22q male and female mice on responses without optogenetic stimulation compared to wild-type controls (Figure 7E-F). This deficit was attenuated with 30hz stimulation in both male and female 22q mice, showing rescued task performance with gamma stimulation. While hit rate was unaffected, both male and female 22q mice optimized their responding rates to non-target stimuli, and within-subjects analysis revealed significant reductions in false alarm rate in both male and female mice with gamma stimulation (Supplementary Figure 8). Finally, to control for off-target effects of optogenetic stimulation, all mice underwent an additional testing session with 5hz stimulation (Supplementary Figure 9). Otherwise, all optogenetic parameters were the same. On this session, both male and female 22q mice still exhibited a deficit in discrimination sensitivity with no effect of 5hz stimulation. Therefore, we can conclude that the rescue of discrimination sensitivity in 22q mice via prefrontal PVN activation was specific to gamma frequency stimulation.

**Figure 7.**
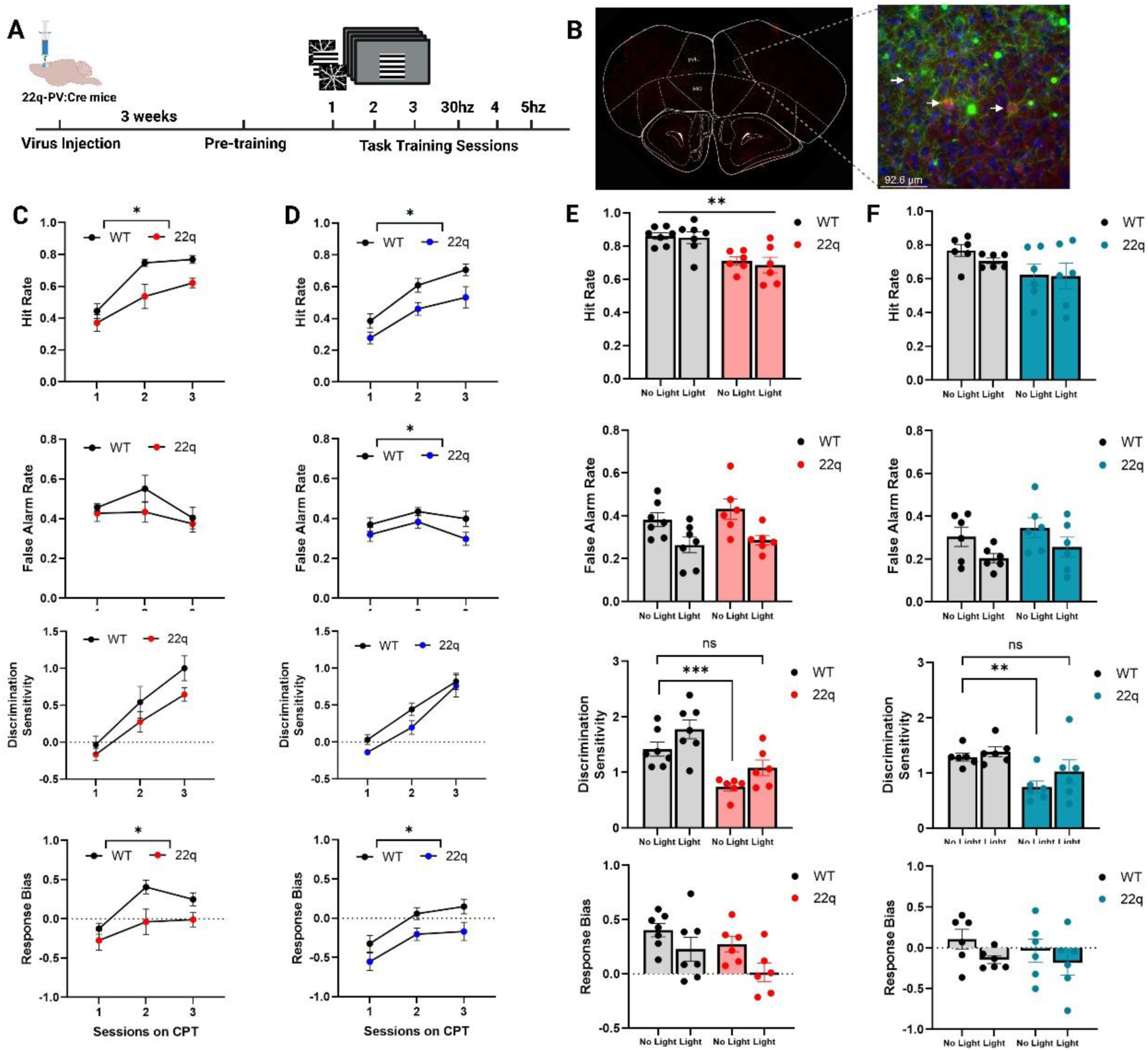
Gamma stimulation of PVNs improves non-target responding and rescues deficits in discrimination sensitivity in a mouse model of 22q11.2 microdeletion syndrome. **A.** The experimental timeline. **B.** (left) Representative coronal slice of the mPFC from a Df(22q):PV-Cre mouse showing co-localization of PV and GFP-tagged ChR2 (right) alongside a higher-powered image (40x). **C.** Male 22q mice (n = 6) display a significant reduction in hit rate (Two-way RM ANOVA: F_1, 11_ = 9.236, p = 0.011) and response bias (F_1, 11_ = 9.599, p = 0.010), compared to WT controls (n = 7). **D.** Female 22q mice (n = 8) display a significant reduction in hit rate (Two-way RM ANOVA: F_1, 12_ = 6.322, p = 0.027), false alarm rate (F_1, 12_ = 5.716, p = 0.034), and response bias (F_1, 12_ = 5.987, p = 0.031), compared to WT controls (n = 6). **E-F.** Gamma frequency stimulation attenuates impairments in discrimination sensitivity in Df(22q):PV-Cre male (left column-red) and female (right column-blue) mice. **E.** Comparison of the effects of stimulating prefrontal PVNs at 30hz in WT and 22q male mice. 30hz stimulation was unable to attenuate the deficit in hit rate between male 22q mice and controls (Two-way ANOVA; main effect of genotype: F_1, 11_ = 14.229, p = 0.003). However, 30hz activation did attenuate the impairment in discrimination sensitivity (Two-way ANOVA; main effect of genotype: F_1, 11_ = 20.863, p < 0.001, and optogenetics: F_1, 11_ = 8.403, p = 0.014). There was a significant reduction in discrimination sensitivity during the “no light” responses (p = 0.0009), but not when 22q mice received 30hz stimulation compared to WT baseline (22q-Light vs. WT-No Light: p = 0.104). **F.** Comparison of the effects of stimulating prefrontal PVNs at 30hz genotypes in female mice. Two 22q mice had to be removed from analysis due to an absence in virus expression (n = 6 each). On this session, we observed no significant differences in hit rate between 22q mice and controls (Two-way ANOVA; no main effect of genotype: F_1, 10_ = 2.692, p = 0.132) Similarly, however, 30hz activation attenuated the impairment in discrimination sensitivity (Two-way ANOVA; main effect of genotype: F_1, 10_ = 7.147, p = 0.023, and optogenetics: F_1, 10_ = 5.182, p = 0.046). Unpaired t-test showed a significant reduction in discrimination sensitivity during the “no light” responses (p = 0.0018), but not when 22q mice received 30hz stimulation compared to WT baseline (22q-Light vs. WT-No Light: p = 0.269). Error bars indicate mean ± SEM. *** p < 0.001, ** p < 0.01, * p ≤ 0.05.

## Discussion

PVNs play a critical role in organizing communication in the cortex, and their regulation of local gamma oscillations makes the functionality of these cells fundamental for cognitive operations. Moreover, the observations of abnormal prefrontal oscillatory activity and PVN deficits in clinical populations has led to interest in their potential as therapeutic targets for cognitive impairments. Here, we show that prefrontal PVNs facilitate performance on a translational task of focused visual attention and were a viable target for improving attention in poorly optimized mice and in a 22qDS mouse model. Importantly, we provide a novel characterization of PVN activity in 22q mutants, demonstrating functional impairments in prefrontal circuitry during task performance. We also observed that PVN signals and their contributions to task performance could be flexible across different task parameters, aligning with their reported involvement in various domains of PFC-dependent cognition.

The identification of neurons in brain regions such as the PFC that display mixed selectivity for multiple task elements suggests that cognition is supported by mechanisms that flexibly facilitate the task at hand [75, 91–95]. Thus, it might be expected that the activity of circuit components such as inhibitory neurons would also dynamically change in response to altered task demands [96–98]. During initial 1NT task training, we observed broad responses of PVNs to stimulus presentation, which may facilitate the tuning of local excitatory neurons to visual stimuli [80, 91, 99], and are in line with previous reports of prefrontal PVN activity during go/no go tasks [2, 80]. In non-pathological mice, PVN signals were strengthened across task learning, specifically when animals were making correct choices for a given stimulus type. Interestingly, although Hits and Correct Rejections are both correct responses, they represent quite distinct situations, differing in stimulus type, response requirement, and association with reward (i.e., only Hits are rewarded). Thus, it may be that the amplification of PVN responses over learning is associated more with general, acquired task rules, rather than specific stimuli or response requirements. It is important to note that our study was limited to population level dynamics, and it is possible that ensembles of prefrontal PVNs may be differentially involved in representations for categories of task rules, such as “go” or “no go” responses [100]. During associative learning, plasticity changes in prefrontal PVN excitability are observed as increases in excitability [101], and PVN modulation has been shown to facilitate learning on paradigms such as fear conditioning [102] and extinction [103]. Moreover, recent studies have shown elevated PVN activity following unexpected rule changes, and the inhibition of prefrontal PVNs disrupts acquiring shifts in rule contingencies [5, 7]. Similarly, updating behavior in response to changing task requirements is also associated with increased mPFC gamma synchrony [6, 17, 18]. Thus, PVNs appear to have a role in the formation of associations and the flexibility to update them when contingencies change. Future research with higher cellular resolution may be able to expand upon whether these cells display flexibility in their representation of task rules.

Although the profiles of PVN activity during stimulus presentation and at response were similar between the 1NT and 4NT tasks, we observed significant changes in PVN signals when task demands were acutely altered by lowering the stimulus duration. The most substantial alterations were observed prior to and following a Hit response, where we observed significant increases in PVN activity peaks. Reducing the stimulus duration during single sessions would be expected to increase the requirement for preparedness prior to stimulus presentation, which appears to be reflected in an increase in PVN activity prior to a correct Hit, and lower PVN activity prior to an incorrect Miss. Interestingly, these data align with previous studies that reported higher PVN activity and gamma band power preceding correct responses during target detection-based attention tasks [10, 11]. Moreover, these results suggest that PVN activity can dynamically shift in response to acute changes in task demands that require behavioral modification, which may reflect similar mechanisms that underlie their involvement in PFC-dependent functions like rule shifting.

Although correctly rejecting a non-target was consistently associated with elevated PVN activity on both versions of the task, we only observed a relationship between PVN activity and task performance on the 4NT task. Specifically, animals that were more proficient at withholding responses to non-targets also displayed significantly higher PVN recruitment when making a Correct Rejection. This association may further aid in explaining the distinct effects of 30hz stimulation on task performance, which facilitated overall discrimination on the 4NT task in low performing mice but was ineffective at modulating behavior in well trained animals on the 1NT task. Thus, it appears that while PVN signals shared similarities across task versions, animals that were poorer at discriminating targets from non-targets benefited from 30hz stimulation when presented with a greater number of non-targets at higher rates (i.e., as is done on the 4NT task). It is important to note, however, that when optogenetic stimulation was provided to 22q mice and wild-type controls on the 1NT task, 30hz stimulation improved false alarm rate in both genotypes. The important difference here is that optogenetic stimulation was provided early in task training well before performance criteria was met. As we observed that PVN signal is enhanced with extensive task training, gamma stimulation early in training may provide artificial circuit optimization and thus improving task performance in non-pathological mice before a ceiling is reached.

## Clinical Relevance

Previous studies have shown that some pharmacological compounds are able to improve attention only in mice that perform more poorly at baseline [84–87]. After confirming that 30hz stimulation of PVNs would successfully entrain local gamma band activity, and 5hz stimulation induced aberrant oscillatory activity, we expected that entraining the local prefrontal circuit to synchronous gamma may improve the processing of incoming visual stimuli in the PFC [14, 81, 104, 105]. In the context of our cognitive paradigm, an enhanced discrimination of targets and non-targets would thereby facilitate task performance, which aligns with our data demonstrating improved discrimination sensitivity with 30hz stimulation. The improvement of task performance, specifically in low performers, suggests that PVNs may be a viable therapeutic target for conditions that exhibit attentional impairments, particularly as a mechanism for restoring synchronous gamma activity.

Deficits in PVNs and cortical synchrony have long been hypothesized to contribute to the cognitive symptoms associated with schizophrenia and are emerging as mechanisms of interest across neuropsychiatric and neurodegenerative diseases [106–108]. Due to the high genetic risk factor for developing psychiatric disorders, the 22qDS model provides a valuable preclinical tool for investigating the neural mechanisms relevant for targeting impairments in cognition. While numerous studies have shown PVN abnormalities in the postmortem PFC tissue of psychiatric patients [20–23, 25–27, 29, 33–35, 109], we provide a real-time functional assessment of PVNs as mutant mice performed a clinically-relevant task of focused visual attention. 22q mice displayed heavily blunted task-related PVN activity and demonstrated an absence of signal strengthening that was present in wild-type mice over task training. A lack of PVN recruitment may be associated with previously described pathology, such as a loss of PV expression, which could contribute to reduced circuit plasticity and inhibit the tuning of PVNs and associated excitatory neurons to task-relevant stimuli over learning [90, 110–112].

Importantly, however, our recordings did show functional features of prefrontal PVN dynamics in 22q mice. The presence of reduced or abnormal function, rather than a global loss of function or cell death, may be a critical feature for the therapeutic potential of targeting PVNs to enhance function in disease. Indeed, we show that stimulating PVNs was effective for improving overall discrimination sensitivity in 22q mice, specifically when it was applied at the gamma frequency. While we were unable to rescue the deficits in target detection, previous work has shown that mutant 22q mice do not exhibit deficits on attention tasks that solely assess identifying and responding to visual targets (e.g., the 5-choice serial reaction time task) [62]. This finding is in agreement with the profiles of clinical populations, who show deficits on CPTs but intact vigilance in the absence of distractors [113]. As a result, the impairment in the 22q mice may be better described as a reduced ability to optimize attention to relevant stimuli. Our findings that gamma stimulation improves cognition in low-performing wild-type and 22q mice suggests an improvement of local circuit processing, and provides support that modulating gamma activity offers a potential therapeutic avenue for psychiatric disorders that display prefrontal cortex dysfunction [5, 6, 11, 114]. Together, these data indicate that PVNs exhibit flexible capabilities, likely to facilitate the dynamic nature of the PFC, and that their disruption has the potential to manifest as dysfunctional behavior across domains of PFC-dependent cognition. The impairment profiles of psychiatric disorders do not present in singular cognitive domains and appear to reflect deficits in prefrontal rhythms and global inefficiencies in synthesizing information towards goal-directed behaviors which rely on organization provided by inhibitory circuits. Thus, targeting prefrontal PVNs as a mechanism for enhancing gamma synchrony may provide a viable treatment avenue for cognitive improvement across a variety of cognitive domains.

## Methods

### Animals

Adult male and female mice aged up to 4 months old were used for all behavioral experiments and mostly bred in-house. All experiments were conducted in compliance with the standards set by the Canadian Council of Animal Care. Prior to experiments, all mice were group housed (2-4 per cage) and maintainded under a reverse light cycle (12hr:12hr) with ad libitum access to food and water. Following optic fiber implant, mice were single housed for the remainder of the experiment. Two weeks prior to touchscreen training, food was restricted to maintain 85-90% of free feeding body weight with ad libitum access to water. Mice were handled daily for one week prior to training and habituated to strawberry milkshake reward (Neilson Strawberry Shake) for three days prior to training. For calcium imaging experiments, homozygous Pvalb-IRES-Cre (PV:Cre: 017320) mice were obtained from Jackson Laboratory and maintained in our mouse colony. For optogenetic experiments, homozygous PV:Cre mice were crossed with Ai40(RCL-ArchT/EGFP)-D (Ai40D) mice or ChR2(H134R)/EYFP (Ai32) mice and maintained in our colony [83]. This generated homozygous PV:Arch and PV:ChR2 offspring will full expression of PVN-selective opsins. Ai40D (021188) and Ai32 (012569) were originally obtained from Jackson Laboratory prior to on-site breeding. C57BL/6-Del(16Dgcr2-Hira)1Tac (22q) mice were obtained from Taconic Biosciences and maintained by crossing hemizygous Df(h22q11.2) mice with C57BL/6 wild-types. For calcium imaging and optogenetics, Df(h22q11.2) mice were crossed with homozygous PV:Cre to generate wild-type and hemizygous 22q offspring with cre expression selective to PVNs. Male and female mice were randomly assigned to experiments and genotypes were confirmed via polymerase chain reaction (PCR) analysis.

### Surgery

Stereotaxic surgery was prior to task training in adult male and female mice. Mice were anesthetized with isuflorance (3% at induction, 2% maintanence in 95% oxygen) and mounted in a stereotaxic frame (David Kopf Instruments). Prior to the operation, subcutaneous Metacam (5mg/kg) was administered for pain control. Body temperature was recorded at the beginning and end of surgery and a heating pad was placed under the mouse to maintain body temperature. An incision was made to expose the skull and bregma and lambda were used as references for coordinate alingment. Bur holes were drilled for virus injection and fiber implant, and two posterior holes for anchoring screws. For calcium imaging experiments, PV:Cre and Df(22q):PV-Cre wild-type and 22q mice were unilaterally injected in the mPFC targeted to the prelimbic cortext (AP: +1.90mm, ML: ±0.30mm, DV: -2.2mm) with 0.5µL of AAV-syn-FLEX-jGCaMP7f-WPRE [78] to selectively target cre-expressing PVNs. Virus was injected using a 10µL Hamilton syringe attached to a microinfusing pump at a rate of 0.1µL/minute. Following the infusion, the syringe was left in place for 5 -10 minutes to prevent backflow before being slowly removed. A single fiber optic probe (400µm core, 0.48 NA, 2.3mm length) was implanted above the injection site and secured to the skull using dental cement (Ketac Cem, 3M). Task training began a minimum of 3 weeks after surgery to allow for adequate biosensor expression.

For optogenetic experiments PV:Arch and PV:ChR2 mice were implanted bilaterally with fiber optic probes (200µm core, 0.22 NA, 2.4mm length) targeted to the prelimbic region of the mPFC (AP: +1.90mm, ML: ±0.30mm, DV: -2.2mm). The coordinates were adjusted to account for a 15° angle which was necessary for attachment to optogenetic patch cables during testing. Mice were given two weeks to recover before food restriction and task training. For optogenetic experiments using the Df(22q):PV-Cre model, wild-type and 22q mice were bilaterally injected with AAV-CAG-DIO-ChR2(H134R) to selectively target cre-expressing PVNs [115] using the same angled coordinates described above. Two fiber optic probes (200µm core, 0.22 NA, 2.4mm length) were lowered into the mPFC above the injection site. Task training began a minimum of 3 weeks after surgery to allow for adequate opsin expression.

### Cognitive tasks

All mice were tested using standard Bussey-Saksida mouse touchscreen chambers (Campden Instruments Ltd., Loughborough, UK) as described in detail elsewhere [116, 117]. The touchscreen was covered with a black plexiglass mask with one row of three response windows. Strawberry milkshake reward was delivered at the back of the chamber, opposite to the screen, in an open reward tray that faciliated mice attached to light sources via patch cables. The task schedules were designed, managed, and events recorded using Whisker Server and ABETTII software (Campden Instruments).

Training on the rCPT closely followed the protocol previously described [63], with minor adjustments. After mice learned to respond to a target image (horizontal lines) that was continuously presented in the center of the screen (2.5s durations), non-target images were added to the session schedule. The first task version, the 1 target, 1 non-target task (1NT), rewarded targets and unrewarded non-target images are continuously presented for a duration of 2.5s at equal probabilities. Mice were required to respond to target images (Hit) and withhold responses to non-target images (Correct Rejection). Incorrect responses included not touching a target image (Miss) or incorrectly touching a non-target image (False Alarm). Image presentation was only briefly halted (5s reward consumption period + variable 2 or 3s ITI) with reward delivery following a correct target Hit. Withholding responses to non-targets (correct response) or targets (incorrect response) simply resulted in the removal of the stimulus. Incorrect responses to a non-target resulted in a correction trial loop, where only non-target images were presented until a response was correctly withheld. The base measures, Hit, Miss, False Alarm, and Correct Rejection are transformed into composite measures using signal detection theory [63, 118, 119]. These measures include:

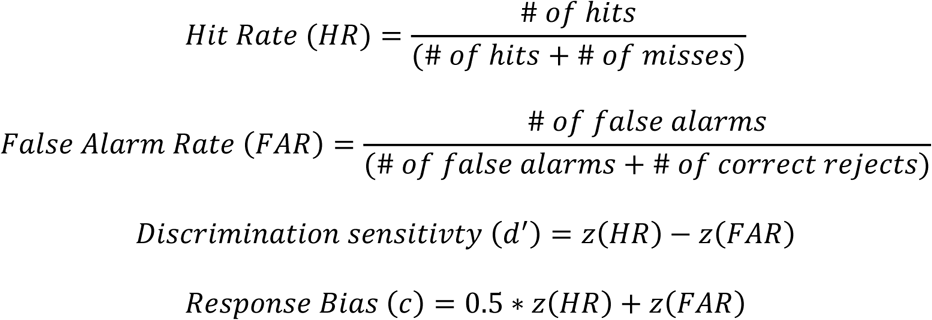

Animals met criterion when they achieved a high performance score of HR > 0.6, FAR < 0.2, d’ > 1.5. Once animals met criterion, they progressed to the 4NT rCPT, which included the same target image from the previous stages, and the addition of four novel non-target image (2x diagonal lines, vertical lines, and circular lines). The probability of image type presentation was not equal for target images, which were presented more infrequently compared to non-targets (33.3% target, 66.6% non-target). Otherwise, all task parameters were the same as the 1NT task.

### Fiber photometry

Fiber photometry experiments were conducted as previously been described [120]. Fiber implants were constructed from optical fiber (0.48 NA, 400 µm core, cut to 2.4 mm; ThorLabs) and secured to stainless steel ferrules (1.25 mm diameter, 6.4 mm long, 440 µm bore; ThorLabs). A patch cable (Doric Lenses) was connected to the fiber optic implanted in the mPFC and real-time fluorescent signals were collected. The photometry system was equipped with a fluorescent mini-cube (Doric Lenses) to transmit two LED light signals at 465nm (sinusoidally modulated at 572hz) and 405nm (modulated at 209hz). Excitation light was delivered at 465nm to stimulate Ca^2+^-dependent GCaMP7f fluorescence. Isosbestic 405nm light was delivered to control for tissue bleaching or movement artifacts, and removed from the 465nm channel prior to analysis. LED power was set at ∼25 μW. Signals were transmitted via the patch cable to the mini-cube, amplified, and focused via an integrated high sensitivity photoreceiver (Doric Lenses). Transistor-transistor logic (TTL) pulses were delivered at session start and continuously at stimulus presentation from ABETTII (Lafayette Instruments). Signals were sampled at 12 kHz and further decimated to 100hz via Doric Studio software V5 (Doric Lenses). The continuous signals were filtered as a function of response type (Hit, Miss, False Alarm, Correct Rejection), averaged across the session, and aligned to task events to assess changes in fluorescence (ΔF/F).

### Electrophysiological recordings

Mice were initially anesthetized under isoflurane (2%) and fixed in a stereotaxic frame with blunt ear bars. An incision was made in the scalp and the tissue overlying the dorsal aspect of the skull was removed with a scalpel. Using a stereotaxic micromanipulator, marks were made on the skull 1.9mm rostral and 0.3mm lateral of Bregma. Subsequently, a craniotomy (1 x 1 mm) was made to expose the cortex overlying the mPFC. A small hole was drilled into the right parietal bone and a stainless-steel screw was inserted to serve as an electrical ground. In preparation for the subsequent extracellular electrophysiological recordings which were conducted under urethane anesthesia, an injection of urethane was given (1.2 g/kg, IP) and the administration isoflurane was terminated. Neural activity was recorded using an optrode that consisted of a 32-channel microelectrode array (H10b; 30 μm site spacing) coupled to a lambda tip optic fiber (100 um core diameter, 0.37 NA, 1mm taper; Cambridge NeuroTech). Using a high-precision stereotaxic manipulator, the optrode was inserted into the cortex through a small slit in the dura, and slowly advanced to a depth of 2.0mm from the cortical surface. Once at the appropriate depth, the optrode was allowed to settle in place for at least 30 min prior to commencing the recordings. The local field potential (LFP) activity was continuously acquired (digitally resampled at 3000 Hz) using a TDT System 3 (Tucker Davis Technologies; TDT) and bandpass filtered online at 1 – 300 hz.

The precise timing of the optogenetic stimulation was controlled via TTL signals sent from the signal processing and acquisition software Synapse (TDT) to a 473nm laser driver (LaserGlow Technologies). The intensity of the blue light emitted from the laser head was controlled by the laser driver, which was coupled to an optic patch cable (200 um core, Doric Lenses) and connected to the optic fiber on the optrode via a plastic sleeve. Optogenetic stimulation consisted of 5ms light pulses that were delivered for 2s at two different frequencies (i.e., 5hz and 30hz). Each stimulation condition (5hz, 30hz, and no light) was presented 100 times in a randomized order, with an intertrial interval of 8s.

### Optogenetic set-up and protocol

Fiber implants were constructed from optical fiber (0.22 NA, 200 µm core, cut to 2.4 mm; ThorLabs) and secured to ceramic ferrules (1.25 mm diameter, 6.4 mm long, 230 µm bore; ThorLabs). Implants attached to two connected optical patch cables (0.66 NA: Plexon) via ceramic mating sleeves, which individually connected to compact LED modules attached to a dual LED commutator above the chamber (PlexBright, Plexon). Light was delivered via a 4-channel optogenetic controller powered by Radiant software (PlexBright, Plexon). Behavioral schedules were coded in ABETTII to synchronize light delivery via Radiant integration to the desired task phase. Light output was measured at the fiber tip via a S140C photodetector connected to a PM100D power meter to confirm adequate light power.

For all optogenetic sessions, light was not delivered until mice completed 10 correct hits to ensure performance was adequately baselined at the beginning of each session. Following this point, all optogenetic stimulation was delivered pseudorandomly throughout the testing session and didn’t occur for more than 3 presentations in a row. For multiple sessions with optogenetic stimulation, a baseline session on the standard task occurred between each one. Optogenetic stimulation was delivered bilaterally during half of stimulus presentations in a pseudorandom manner (Session: Max 100 hits, 45-minute duration). Light delivery occurred until a response was made, or the stimulus was removed from the screen. For optogenetic inhibition experiments, green light (550nm, ̴ 2.5mW, constant) was delivered starting at the final second of the ITI and through the stimulus presentation (max 3.5s). Additional testing sessions included delivering inhibitory light during the ITI period (max 3s) or the stimulus presentation period (max 2.5s). For optogenetic activation experiments, excitation light (465nm, 5ms pulses, ̴ 1mW) was similarly delivered starting at the final second of the ITI and through the stimulus presentation. Frequency specific light patterns were coded in Radiant software and were otherwise delivered identically. Testing sessions were counterbalanced with 30hz or 5hz frequency stimulation to prevent testing order effects. During LFP experiments, the same optical parameters were used to ensure this protocol was effective at inducing corresponding oscillatory activity.

### Histology

Mice were anesthetized with a mixture of ketamine and xylazine prior to transcardial perfusion with PBS, followed by 4% paraformaldehye (PFA). Brains were extracted and postfixed overnight in 4% PFA at 4°C, before being transferred to PBS with 30% sucrose solution for cryoprotection. Brains were coronally sliced in the mPFC region at 40µm using a vibratome (Leica Biosystems). For immunostaining, slices were mounted to slides, incubated in Tris-buffered saline (TBS) with 1.2% Triton X-100 for 20 min, and blocked with TBS containing 5% normal goat serum (NGS) at room temperature for one hour. Sections were incubated overnight at 4°C with rabbit monoclonal anti-parvalbumin antibody (1:50; Abcam 181086) in TBS with 0.2% Triton X-100 and 2% NGS. Next, sections were incubated in TBS with 0.2% Triton X-100 and 2% NGSwith secondary antibody Alexa Fluor 594-conjugated goat anti-rabbit (Invitrogen A11012) for one hour before an overnight incubation in primary anti-GFP polyclonal 488-conjugated rabbit (1:1000; Abcam 21311). Lastly, sections were coverslipped with Vectashield with DAPI (MJS BioLynx, VECTH1200) for visualization. Virus expression and fiber optic placement was confirmed using a Thunder imaging microscope (Leica Microsystems) and representative images were processed using ImageJ.

### Fiber photometry analysis

To assess peak GCaMP7f signals, normalized changes in fluorescence (ΔF/F) were calculated as: ΔF/F = [465 nm signal – (fitted 405 nm signal)]/(fitted 405 nm signal). Calcium signals were baselined to the initial second of the ITI based on assessment of signal variance across non-event related time periods. To assess PVN activity during stimulus presentation, ΔF/F was calculated as: Δ in signal = (ΔF/F – μbaseline)/σbaseline, where μ baseline is the mean of values from baseline period (averaged signal collected 1s before stimulus onset) and σbaseline is the standard deviation of values from baseline period [120]. Raw trial-by-trial data was analyzed using a custom-written Python script (Palmer, 2022; Available at: https://doi.org/10.5281/zenodo.7577053). GCaMP7f signals were aligned to stimulus presentation or time of response and the data output was organized for area under the curve (AUC) and max signal peak analysis at specific time points using OriginLab software. Signals were analyzed during the 0.5s after stimulus presentation or 0.5s prior to response periods, to assess activity at temporally specific events. Sampling also occurred during the final second of the ITI immediately prior to stimulus presentation, following a non-rewarded response (2s period), and during reward collection (2s period). There were no sex differences observed in behavioral performance or photometry signal, so male and female mice were combined for analysis.

### LFP analysis

Custom Matlab scripts and functions from the FieldTrip toolbox [121] were used to quantify the consequence of the frequency-specific optogenetic stimulation on the neural activity recorded within the mPFC. The LFP values were down sampled to 1000 Hz, converted to µV, and averaged across the trials for the different optogenetic stimulation conditions (5hz, 30hz and no light). Each trial was subjected to a time-frequency decomposition using the ‘*mtmconvol*’ method with a Hanning window taper. Evoked power (EP) and intertrial coherence (ITC) were then calculated from the complex value generated for each frequency of interest (i.e., 0 to 60hz in 0.5hz steps) from the beginning to the end of the trial (i.e., 0 to 8s) using a 200ms window centered on 1ms steps. EP was calculated by averaging the complex values across all trials and then squaring the magnitudes, whereas the ITC was calculated by dividing each complex value by its magnitude. Both EP and ITC were baseline corrected using a time window of a -600 to -100ms with respect to stimulus onset at each oscillatory frequency of interest. Both metrics were further quantified within the optogenetic stimulation period (i.e., 0 to 2s), by calculating mean EP and ITC within three frequency bins of interest: delta/theta (Δ/Ɵ), 1 – 9hz; low-gamma (low-γ), 26 – 34hz; high-gamma (high-γ), 51 – 59hz, which were then averaged across mice for each stimulation condition (5hz, 30hz and no light). Lastly, the overall oscillatory profile was compared between the optogenetic stimulation conditions by computing the scaled power of the oscillations. The LFP values during the optogenetic simulation were subjected to a time-frequency decomposition via a Fast-Fourier Transformation (FFT) that utilized a Hanning window taper. The magnitudes of the resulting complex values were squared to yield power values, and then were averaged across trials to create a single power spectrum. Each power spectrum was normalized by its overall mean power, thereby yielding a scaled power [122–125]. Similar to the EP and ITC values, further quantification was performed by calculating the mean scaled power within the three frequency bins of interest (i.e., delta/theta, low-gamma and high-gamma).

### Behavioral data and statistical analysis

Behavioral data was extracted from ABETTII to analyze performance measures across training and during optical testing. Performance measures were transformed to composite measures hit rate (HR), false alarm rate (FAR), discrimination sensivivity (d’), and response bias (c) before statistical analysis was performed. A custom R script was used to analyze behavioral responses with vs. without optogenetic stimulation. Latency measures, including correct and incorrect choice latency describe the time in seconds to respond to the screen following the presentation of the original and novel stimuli during the choice phase. Reward collection latency was the time in seconds it took a mouse to return to the reward tray following a correct response. Graphs were generated using GraphPad Prism, and data is shown as the mean with error bars representating standard error of the mean (SEM). Group comparisons were done using repeated measures ANOVA, and included factors of genotype (when assessing the 22qDS model), sex, and condition when applicable. Conditions included optogenetic manipulations and photometry signals across various task phases. Tukey’s post-hoc test was used following ANOVAs when analyzing multiple comparisons. Post-hoc paired or unpaired student’s t-tests were used to make single variable comparisons. Data distribution was assumed to be normal but this was not formally tested. All data were checked by the Shapiro–Wilk test and Leven’s test for normality and homogeneity of variance, respectively. Violation of sphericity assessed by Mauchly’s test was corrected using the Greenhouse–Geisser method. No statistical methods were used to predetermine sample size, but our sample sizes are similar to previous investigations in the field using similar techniques [5–7, 90, 126]. All statistical analyses were conducted using JASP. Statistical analysis yeilding p ≥ 0.05 is considered not significant and denoted by the lack of an asterick. p = *, < 0.05; **, < 0.01; ***, < 0.001; ****, < 0.0001.

## Acknowledgments

This research was supported by through the Canada First Research Excellence Fund (CFREF), Brain Canada, the Canadian Institutes of Health Research (CHIR PJT 426966), the Natural Sciences and Engineering Research Council (NSERC RGPIN-2019-06102 RGPIN-2019-06087), the Canada Foundation for Innovation, and the Ontario Research Fund. LMS is the Canada Research Chair in Translational Cognitive Neuroscience and TJB is the Western Research Chair in Behavioural Neuroscience. Figures for the manuscript were created using Biorender (https://BioRender.com).

## Author Contributions

The project was conceived by T.D.D, T.J.B, L.M.S., and D.P. and supervised by D.P., T.J.B. and L.M.S. T.D.D performed the behavioral experiments, analyzed the data, generated the figures, and wrote the manuscript. T.D.D., T.J.B, and L.M.S edited the manuscript. A.L.S. conducted the electrophysiology experiments, analyzed the data, generated Figure 4, and wrote the electrophysiology results and methods sections. D.P. wrote custom photometry and optogenetic analysis scripts and provided training and supervision. S.H. assisted in testing mice included in Supplementary Figure 5. T.D.D. and M.M.F.M. processed the brain tissue for histology. M.M.F.M. performed the immunohistochemistry and generated the images. B.L.A. provided insight throughout the project and supervised the electrophysiology experiments.

## Competing interests

T.J.B. and L.M.S. have established a series of targeted cognitive tests for animals, administered via touchscreen within a custom environment known as the “Bussey-Saksida touchscreen chamber”. Cambridge Enterprise, the technology transfer office of the University of Cambridge, supported commercialisation of the Bussey-Saksida chamber, culminating in a license to Campden Instruments. Any financial compensation received from commercialisation of the technology is fully invested in further touchscreen development and/or maintenance. T.D., D.P., A.L.S., S.H., M.M.F.M., B.L.A. declare no competing interests.

## Data Availability

Data is available upon request. Additionally, the datasets generated and analysed in this study will be freely accessible in the Mousebytes repository, https://mousebytes.ca/home.

## References

1. Murray, A.J., et al., Parvalbumin-positive interneurons of the prefrontal cortex support working memory and cognitive flexibility. Scientific reports, 2015. 5(1): p. 16778.

2. Kamigaki, T. and Y. Dan, Delay activity of specific prefrontal interneuron subtypes modulates memory-guided behavior. Nature neuroscience, 2017. 20(6): p. 854–863.

3. Nguyen, R., et al., Cholecystokinin-expressing interneurons of the medial prefrontal cortex mediate working memory retrieval. Journal of Neuroscience, 2020. 40(11): p. 2314–2331.

4. Chamberlin, L.A., et al., Pharmacogenetic activation of parvalbumin interneurons in the prefrontal cortex rescues cognitive deficits induced by adolescent MK801 administration. Neuropsychopharmacology, 2023. 48(9): p. 1267–1276.

5. Cho, K.K., et al., Cross-hemispheric gamma synchrony between prefrontal parvalbumin interneurons supports behavioral adaptation during rule shift learning. Nature neuroscience, 2020. 23(7): p. 892–902.

6. Cho, K.K., et al., Gamma rhythms link prefrontal interneuron dysfunction with cognitive inflexibility in Dlx5/6+/− mice. Neuron, 2015. 85(6): p. 1332–1343.

7. Cho, K.K., et al., Long-range inhibition synchronizes and updates prefrontal task activity. Nature, 2023. 617(7961): p. 548–554.

8. Toader, O., et al., Suppression of parvalbumin interneuron activity in the prefrontal cortex recapitulates features of impaired excitatory/inhibitory balance and sensory processing in schizophrenia. Schizophrenia bulletin, 2020. 46(4): p. 981–989.

9. Patrono, E., et al., The role of optogenetic stimulations of parvalbumin-positive interneurons in the prefrontal cortex and the ventral hippocampus on an acute MK-801 model of schizophrenia-like cognitive inflexibility. Schizophrenia Research, 2023. 252: p. 198–205.

10. Kim, H., et al., Prefrontal parvalbumin neurons in control of attention. Cell, 2016. 164(1): p. 208–218.

11. Ferguson, B., C. Glick, and J.R. Huguenard, Prefrontal PV interneurons facilitate attention and are linked to attentional dysfunction in a mouse model of absence epilepsy. Elife, 2023. 12: p. e78349.

12. Sohal, V.S., et al., Parvalbumin neurons and gamma rhythms enhance cortical circuit performance. Nature, 2009. 459(7247): p. 698–702.

13. Cardin, J.A., et al., Driving fast-spiking cells induces gamma rhythm and controls sensory responses. Nature, 2009. 459(7247): p. 663–667.

14. Gregoriou, G.G., et al., High-frequency, long-range coupling between prefrontal and visual cortex during attention. science, 2009. 324(5931): p. 1207–1210.

15. Buzsáki, G. and X.-J. Wang, Mechanisms of gamma oscillations. Annual review of neuroscience, 2012. 35: p. 203–225.

16. Tamura, M., et al., Hippocampal-prefrontal theta-gamma coupling during performance of a spatial working memory task. Nature communications, 2017. 8(1): p. 2182.

17. Karlsson, M.P., D.G. Tervo, and A.Y. Karpova, Network resets in medial prefrontal cortex mark the onset of behavioral uncertainty. Science, 2012. 338(6103): p. 135–139.

18. Marton, T.F., et al., Roles of prefrontal cortex and mediodorsal thalamus in task engagement and behavioral flexibility. Journal of Neuroscience, 2018. 38(10): p. 2569–2578.

19. Beasley, C.L. and G.P. Reynolds, Parvalbumin-immunoreactive neurons are reduced in the prefrontal cortex of schizophrenics. Schizophrenia research, 1997. 24(3): p. 349–355.

20. Beneyto, M. and D.A. Lewis, Insights into the neurodevelopmental origin of schizophrenia from postmortem studies of prefrontal cortical circuitry. International Journal of Developmental Neuroscience, 2011. 29(3): p. 295–304.

21. Chung, D.W., K.N. Fish, and D.A. Lewis, Pathological basis for deficient excitatory drive to cortical parvalbumin interneurons in schizophrenia. American Journal of Psychiatry, 2016. 173(11): p. 1131–1139.

22. Enwright III, J.F., et al., Transcriptome alterations of prefrontal cortical parvalbumin neurons in schizophrenia. Molecular psychiatry, 2018. 23(7): p. 1606–1613.

23. Glausier, J., K. Fish, and D. Lewis, Altered parvalbumin basket cell inputs in the dorsolateral prefrontal cortex of schizophrenia subjects. Molecular psychiatry, 2014. 19(1): p. 30–36.

24. Gonzalez-Burgos, G., R.Y. Cho, and D.A. Lewis, Alterations in cortical network oscillations and parvalbumin neurons in schizophrenia. Biological psychiatry, 2015. 77(12): p. 1031–1040.

25. Hashimoto, T., et al., Gene expression deficits in a subclass of GABA neurons in the prefrontal cortex of subjects with schizophrenia. Journal of Neuroscience, 2003. 23(15): p. 6315–6326.

26. Hoftman, G.D., D. Datta, and D.A. Lewis, Layer 3 excitatory and inhibitory circuitry in the prefrontal cortex: developmental trajectories and alterations in schizophrenia. Biological psychiatry, 2017. 81(10): p. 862–873.

27. Lewis, D.A., et al., Lamina-specific deficits in parvalbumin-immunoreactive varicosities in the prefrontal cortex of subjects with schizophrenia: evidence for fewer projections from the thalamus. American Journal of Psychiatry, 2001. 158(9): p. 1411–1422.

28. Lewis, D.A., et al., Cortical parvalbumin interneurons and cognitive dysfunction in schizophrenia. Trends in neurosciences, 2012. 35(1): p. 57–67.

29. Kaar, S.J., et al., Pre-frontal parvalbumin interneurons in schizophrenia: a meta-analysis of post-mortem studies. Journal of Neural Transmission, 2019. 126: p. 1637–1651.

30. Senkowski, D. and J. Gallinat, Dysfunctional prefrontal gamma-band oscillations reflect working memory and other cognitive deficits in schizophrenia. Biological psychiatry, 2015. 77(12): p. 1010–1019.

31. Uhlhaas, P.J. and W. Singer, Abnormal neural oscillations and synchrony in schizophrenia. Nature reviews neuroscience, 2010. 11(2): p. 100–113.

32. Haenschel, C., et al., Cortical oscillatory activity is critical for working memory as revealed by deficits in early-onset schizophrenia. Journal of Neuroscience, 2009. 29(30): p. 9481–9489.

33. Toker, L., et al., Transcriptomic evidence for alterations in astrocytes and parvalbumin interneurons in subjects with bipolar disorder and schizophrenia. Biological psychiatry, 2018. 84(11): p. 787–796.

34. Dienel, S.J., K.N. Fish, and D.A. Lewis, The nature of prefrontal cortical GABA neuron alterations in schizophrenia: markedly lower somatostatin and parvalbumin gene expression without missing neurons. American Journal of Psychiatry, 2023. 180(7): p. 495–507.

35. Enwright, J.F., et al., Reduced labeling of parvalbumin neurons and perineuronal nets in the dorsolateral prefrontal cortex of subjects with schizophrenia. Neuropsychopharmacology, 2016. 41(9): p. 2206–2214.

36. Arai, H., et al., Loss of parvalbumin-immunoreactive neurones from cortex in Alzheimer-type dementia. Brain research, 1987. 418(1): p. 164–169.

37. Satoh, J., et al., Parvalbumin-immunoreactive neurons in the human central nervous system are decreased in Alzheimer’s disease. Acta neuropathologica, 1991. 81: p. 388–395.

38. Mikkonen, M., et al., Subfield-and layer-specific changes in parvalbumin, calretinin and calbindin-D28K immunoreactivity in the entorhinal cortex in Alzheimer’s disease. Neuroscience, 1999. 92(2): p. 515–532.

39. Verret, L., et al., Inhibitory interneuron deficit links altered network activity and cognitive dysfunction in Alzheimer model. Cell, 2012. 149(3): p. 708–721.

40. Iaccarino, H.F., et al., Gamma frequency entrainment attenuates amyloid load and modifies microglia. Nature, 2016. 540(7632): p. 230–235.

41. Lanoue, A.C., G.J. Blatt, and J.-J. Soghomonian, Decreased parvalbumin mRNA expression in dorsolateral prefrontal cortex in Parkinson′ s disease. Brain research, 2013. 1531: p. 37–47.

42. Lilascharoen, V., et al., Divergent pallidal pathways underlying distinct Parkinsonian behavioral deficits. Nature neuroscience, 2021. 24(4): p. 504–515.

43. Reiner, A., et al., Striatal parvalbuminergic neurons are lost in Huntington’s disease: implications for dystonia. Movement Disorders, 2013. 28(12): p. 1691–1699.

44. Kim, E.H., et al., Cortical interneuron loss and symptom heterogeneity in Huntington disease. Annals of neurology, 2014. 75(5): p. 717–727.

45. Edelmann, L., R.K. Pandita, and B.E. Morrow, Low-copy repeats mediate the common 3-Mb deletion in patients with velo-cardio-facial syndrome. The American Journal of Human Genetics, 1999. 64(4): p. 1076–1086.

46. Karayiorgou, M., et al., Schizophrenia susceptibility associated with interstitial deletions of chromosome 22q11. Proceedings of the National Academy of Sciences, 1995. 92(17): p. 7612–7616.

47. Shaikh, T.H., et al., Chromosome 22-specific low copy repeats and the 22q11. 2 deletion syndrome: genomic organization and deletion endpoint analysis. Human molecular genetics, 2000. 9(4): p. 489–501.

48. Maynard, T., et al., A comprehensive analysis of 22q11 gene expression in the developing and adult brain. Proceedings of the National Academy of Sciences, 2003. 100(24): p. 14433–14438.

49. Karayiorgou, M., T.J. Simon, and J.A. Gogos, 22q11. 2 microdeletions: linking DNA structural variation to brain dysfunction and schizophrenia. Nature Reviews Neuroscience, 2010. 11(6): p. 402–416.

50. Schneider, M., et al., Psychiatric disorders from childhood to adulthood in 22q11.2 deletion syndrome: results from the International Consortium on Brain and Behavior in 22q11.2 Deletion Syndrome. Am J Psychiatry, 2014. 171(6): p. 627–39.

51. Consortium, W.T.C.C., et al., Microduplications of 16p11.2 are associated with schizophrenia. Nature Genetics, 2009. 41(11): p. 1223–1227.

52. Simon, T.J., et al., A multilevel analysis of cognitive dysfunction and psychopathology associated with chromosome 22q11. 2 deletion syndrome in children. Development and psychopathology, 2005. 17(3): p. 753–784.

53. Sobin, C., et al., Networks of attention in children with the 22q11 deletion syndrome. Developmental neuropsychology, 2004. 26(2): p. 611–626.

54. Bish, J.P., et al., Maladaptive conflict monitoring as evidence for executive dysfunction in children with chromosome 22q11. 2 deletion syndrome. Developmental science, 2005. 8(1): p. 36–43.

55. Stoddard, J., L. Beckett, and T.J. Simon, Atypical development of the executive attention network in children with chromosome 22q11. 2 deletion syndrome. Journal of Neurodevelopmental Disorders, 2011. 3(1): p. 76–85.

56. Larsen, K.M., et al., 22q11. 2 deletion syndrome is associated with impaired auditory steady-state gamma response. Schizophrenia Bulletin, 2018. 44(2): p. 388–397.

57. Mancini, V., et al., Oscillatory neural signatures of visual perception across developmental stages in individuals with 22q11. 2 deletion syndrome. Biological Psychiatry, 2022. 92(5): p. 407–418.

58. Mancini, V., et al., Aberrant developmental patterns of gamma-band response and long-range communication disruption in youths with 22q11. 2 deletion syndrome. American Journal of Psychiatry, 2022. 179(3): p. 204–215.

59. Tripathi, A., et al., Cognition-and circuit-based dysfunction in a mouse model of 22q11. 2 microdeletion syndrome: effects of stress. Translational Psychiatry, 2020. 10(1): p. 41.

60. Al-Absi, A.-R., et al., Df (h22q11)/+ mouse model exhibits reduced binding levels of GABAA receptors and structural and functional dysregulation in the inhibitory and excitatory networks of hippocampus. Molecular and Cellular Neuroscience, 2022. 122: p. 103769.

61. Nilsson, S.R., et al., Assessing the cognitive translational potential of a mouse model of the 22q11. 2 microdeletion syndrome. Cerebral cortex, 2016. 26(10): p. 3991–4003.

62. Nilsson, S.R., et al., Continuous performance test impairment in a 22q11. 2 microdeletion mouse model: improvement by amphetamine. Translational Psychiatry, 2018. 8(1): p. 247.

63. Kim, C.H., et al., The continuous performance test (rCPT) for mice: a novel operant touchscreen test of attentional function. Psychopharmacology (Berl), 2015. 232(21-22): p. 3947–66.

64. Mar, A.C., et al., MAM-E17 rat model impairments on a novel continuous performance task: effects of potential cognitive enhancing drugs. Psychopharmacology, 2017. 234: p. 2837–2857.

65. Nestor, P.G., et al., Measurement of visual sustained attention in schizophrenia using signal detection analysis and a newly developed computerized CPT task. Schizophr Res, 1990. 3(5-6): p. 329–32.

66. Cornblatt, B.A. and J.G. Keilp, Impaired attention, genetics, and the pathophysiology of schizophrenia. Schizophrenia bulletin, 1994. 20(1): p. 31–46.

67. Snitz, B.E., A.W. MacDonald III, and C.S. Carter, Cognitive deficits in unaffected first-degree relatives of schizophrenia patients: a meta-analytic review of putative endophenotypes. 2006.

68. Lewandowski, K.E., et al., Schizophrenic-like neurocognitive deficits in children and adolescents with 22q11 deletion syndrome. American Journal of Medical Genetics Part B: Neuropsychiatric Genetics, 2007. 144(1): p. 27–36.

69. Hooper, S.R., et al., A longitudinal examination of the psychoeducational, neurocognitive, and psychiatric functioning in children with 22q11. 2 deletion syndrome. Research in developmental disabilities, 2013. 34(5): p. 1758–1769.

70. Nuechterlein, K.H., Vigilance in schizophrenia and related disorders. 1991.

71. Miller, E.K. and J.D. Cohen, An integrative theory of prefrontal cortex function. Annu Rev Neurosci, 2001. 24: p. 167–202.

72. Gaffan, D., Against memory systems. Philosophical Transactions of the Royal Society of London. Series B: Biological Sciences, 2002. 357(1424): p. 1111–1121.

73. Bussey, T.J. and L.M. Saksida, Object memory and perception in the medial temporal lobe: an alternative approach. Current opinion in neurobiology, 2005. 15(6): p. 730–737.

74. Forwood, S.E., et al., Multiple cognitive abilities from a single cortical algorithm. Journal of Cognitive Neuroscience, 2012. 24(9): p. 1807–1825.

75. Panichello, M.F. and T.J. Buschman, Shared mechanisms underlie the control of working memory and attention. Nature, 2021. 592(7855): p. 601–605.

76. Didriksen, M., et al., Persistent gating deficit and increased sensitivity to NMDA receptor antagonism after puberty in a new mouse model of the human 22q11. 2 microdeletion syndrome: a study in male mice. Journal of Psychiatry and Neuroscience, 2017. 42(1): p. 48–58.

77. Hvoslef-Eide, M., et al., Effects of anterior cingulate cortex lesions on a continuous performance task for mice. Brain Neurosci Adv, 2018. 2.

78. Dana, H., et al., High-performance calcium sensors for imaging activity in neuronal populations and microcompartments. Nature methods, 2019. 16(7): p. 649–657.

79. Insel, N. and C.A. Barnes, Differential activation of fast-spiking and regular-firing neuron populations during movement and reward in the dorsal medial frontal cortex. Cerebral Cortex, 2015. 25(9): p. 2631–2647.

80. Pinto, L. and Y. Dan, Cell-type-specific activity in prefrontal cortex during goal-directed behavior. Neuron, 2015. 87(2): p. 437–450.

81. Kim, D., et al., Distinct roles of parvalbumin-and somatostatin-expressing interneurons in working memory. Neuron, 2016. 92(4): p. 902–915.

82. Jeong, H., et al., Distinct roles of parvalbumin-and somatostatin-expressing neurons in flexible representation of task variables in the prefrontal cortex. Progress in Neurobiology, 2020. 187: p. 101773.

83. Madisen, L., et al., A toolbox of Cre-dependent optogenetic transgenic mice for light-induced activation and silencing. Nature neuroscience, 2012. 15(5): p. 793–802.

84. Robinson, E.S., Blockade of noradrenaline re-uptake sites improves accuracy and impulse control in rats performing a five-choice serial reaction time tasks. Psychopharmacology, 2012. 219: p. 303–312.

85. Tomlinson, A., et al., Pay attention to impulsivity: modelling low attentive and high impulsive subtypes of adult ADHD in the 5-choice continuous performance task (5C-CPT) in female rats. Eur Neuropsychopharmacol, 2014. 24(8): p. 1371–80.

86. Caballero-Puntiverio, M., et al., Effects of amphetamine and methylphenidate on attentional performance and impulsivity in the mouse 5-Choice Serial Reaction Time Task. Journal of Psychopharmacology, 2017. 31(2): p. 272–283.

87. Caballero-Puntiverio, M., et al., Effect of ADHD medication in male C57BL/6J mice performing the rodent Continuous Performance Test. Psychopharmacology, 2019. 236: p. 1839–1851.

88. Liu, S.K., et al., Deficits in sustained attention in schizophrenia and affective disorders: stable versus state-dependent markers. American Journal of Psychiatry, 2002. 159(6): p. 975–982.

89. Filice, F., et al., Reduction in parvalbumin expression not loss of the parvalbumin-expressing GABA interneuron subpopulation in genetic parvalbumin and shank mouse models of autism. Molecular brain, 2016. 9: p. 1–17.

90. Mukherjee, A., et al., Long-lasting rescue of network and cognitive dysfunction in a genetic schizophrenia model. Cell, 2019. 178(6): p. 1387–1402. e14.

91. Asaad, W.F., G. Rainer, and E.K. Miller, Neural activity in the primate prefrontal cortex during associative learning. Neuron, 1998. 21(6): p. 1399–1407.

92. Mansouri, F.A., K. Matsumoto, and K. Tanaka, Prefrontal cell activities related to monkeys’ success and failure in adapting to rule changes in a Wisconsin Card Sorting Test analog. Journal of Neuroscience, 2006. 26(10): p. 2745–2756.

93. Hussar, C.R. and T. Pasternak, Flexibility of sensory representations in prefrontal cortex depends on cell type. Neuron, 2009. 64(5): p. 730–743.

94. Rigotti, M., et al., The importance of mixed selectivity in complex cognitive tasks. Nature, 2013. 497(7451): p. 585–590.

95. Parthasarathy, A., et al., Mixed selectivity morphs population codes in prefrontal cortex. Nature neuroscience, 2017. 20(12): p. 1770–1779.

96. Wilent, W.B. and D. Contreras, Dynamics of excitation and inhibition underlying stimulus selectivity in rat somatosensory cortex. Nature neuroscience, 2005. 8(10): p. 1364–1370.

97. Yoshimura, Y. and E.M. Callaway, Fine-scale specificity of cortical networks depends on inhibitory cell type and connectivity. Nature neuroscience, 2005. 8(11): p. 1552–1559.

98. Roach, J.P., A.K. Churchland, and T.A. Engel, Choice selective inhibition drives stability and competition in decision circuits. Nature communications, 2023. 14(1): p. 147.

99. Lee, S.-H., et al., Activation of specific interneurons improves V1 feature selectivity and visual perception. Nature, 2012. 488(7411): p. 379–383.

100. Reinert, S., et al., Mouse prefrontal cortex represents learned rules for categorization. Nature, 2021. 593(7859): p. 411–417.

101. Xu, Y., et al., ErbB4 in parvalbumin-positive interneurons mediates proactive interference in olfactory associative reversal learning. Neuropsychopharmacology, 2022. 47(7): p. 1292–1303.

102. Courtin, J., et al., Prefrontal parvalbumin interneurons shape neuronal activity to drive fear expression. Nature, 2014. 505(7481): p. 92–96.

103. Sparta, D.R., et al., Activation of prefrontal cortical parvalbumin interneurons facilitates extinction of reward-seeking behavior. Journal of Neuroscience, 2014. 34(10): p. 3699–3705.

104. Salinas, E. and T.J. Sejnowski, Correlated neuronal activity and the flow of neural information. Nature reviews neuroscience, 2001. 2(8): p. 539–550.

105. Gregoriou, G.G., S. Paneri, and P. Sapountzis, Oscillatory synchrony as a mechanism of attentional processing. Brain research, 2015. 1626: p. 165–182.

106. Santos-Silva, T., et al., Prefrontal and hippocampal parvalbumin interneurons in animal models for schizophrenia: a systematic review and meta-analysis. Schizophrenia Bulletin, 2024. 50(1): p. 210–223.

107. Hijazi, S., A.B. Smit, and R.E. van Kesteren, Fast-spiking parvalbumin-positive interneurons in brain physiology and Alzheimer’s disease. Molecular Psychiatry, 2023. 28(12): p. 4954–4967.

108. Terstege, D.J. and J.R. Epp, Parvalbumin as a sex-specific target in Alzheimer’s disease research, A mini-review. Neuroscience & Biobehavioral Reviews, 2023: p. 105370.

109. Fung, S.J., et al., Schizophrenia and bipolar disorder show both common and distinct changes in cortical interneuron markers. Schizophrenia research, 2014. 155(1-3): p. 26–30.

110. Caillard, O., et al., Role of the calcium-binding protein parvalbumin in short-term synaptic plasticity. Proceedings of the National Academy of Sciences, 2000. 97(24): p. 13372–13377.

111. Orduz, D., et al., Parvalbumin tunes spike-timing and efferent short-term plasticity in striatal fast spiking interneurons. The Journal of physiology, 2013. 591(13): p. 3215–3232.

112. Wöhr, M., et al., Lack of parvalbumin in mice leads to behavioral deficits relevant to all human autism core symptoms and related neural morphofunctional abnormalities. Translational psychiatry, 2015. 5(3): p. e525–e525.

113. Demeter, E., et al., Increased distractor vulnerability but preserved vigilance in patients with schizophrenia: evidence from a translational Sustained Attention Task. Schizophr Res, 2013. 144(1-3): p. 136–41.

114. Cao, W., et al., Gamma oscillation dysfunction in mPFC leads to social deficits in neuroligin 3 R451C knockin mice. Neuron, 2018. 97(6): p. 1253–1260. e7.

115. Rajendran, P.S., et al., Identification of peripheral neural circuits that regulate heart rate using optogenetic and viral vector strategies. Nature communications, 2019. 10(1): p. 1944.

116. Horner, A.E., et al., The touchscreen operant platform for testing learning and memory in rats and mice. Nature Protocols, 2013. 8(10): p. 1961–1984.

117. Oomen, C.A., et al., The touchscreen operant platform for testing working memory and pattern separation in rats and mice. Nature Protocols, 2013. 8(10): p. 2006–2021.

118. Green, D.M. and J.A. Swets, Signal detection theory and psychophysics. Vol. 1. 1966: Wiley New York.

119. Frey, P.W. and J.A. Colliver, Sensitivity and responsivity measures for discrimination learning. Learning and Motivation, 1973. 4(3): p. 327–342.

120. Skirzewski, M., et al., Continuous cholinergic-dopaminergic updating in the nucleus accumbens underlies approaches to reward-predicting cues. Nature Communications, 2022. 13(1): p. 7924.

121. Oostenveld, R., et al., FieldTrip: open source software for advanced analysis of MEG, EEG, and invasive electrophysiological data. Computational intelligence and neuroscience, 2011. 2011(1): p. 156869.

122. Hayes, S.H., et al., Uncovering the contribution of enhanced central gain and altered cortical oscillations to tinnitus generation. Progress in Neurobiology, 2021. 196: p. 101893.

123. Wieczerzak, K.B., et al., Differential plasticity in auditory and prefrontal cortices, and cognitive-behavioral deficits following noise-induced hearing loss. Neuroscience, 2021. 455: p. 1–18.

124. Weisz, N., K. Dohrmann, and T. Elbert, The relevance of spontaneous activity for the coding of the tinnitus sensation. Progress in brain research, 2007. 166: p. 61–70.

125. Weisz, N., et al., Tinnitus perception and distress is related to abnormal spontaneous brain activity as measured by magnetoencephalography. PLoS medicine, 2005. 2(6): p. e153.

126. Norman, K.J., et al., Chemogenetic suppression of anterior cingulate cortical neurons projecting to the visual cortex disrupts attentional behavior in mice. Neuropsychopharmacology Reports, 2021. 41(2): p. 207–214.

