## Supplementary Figures for "Prefrontal parvalbumin neurons as a target for enhancing cognition in non-pathological and 22q11.2 microdeletion syndrome mice": Manuscript Supplementary Information .pdf

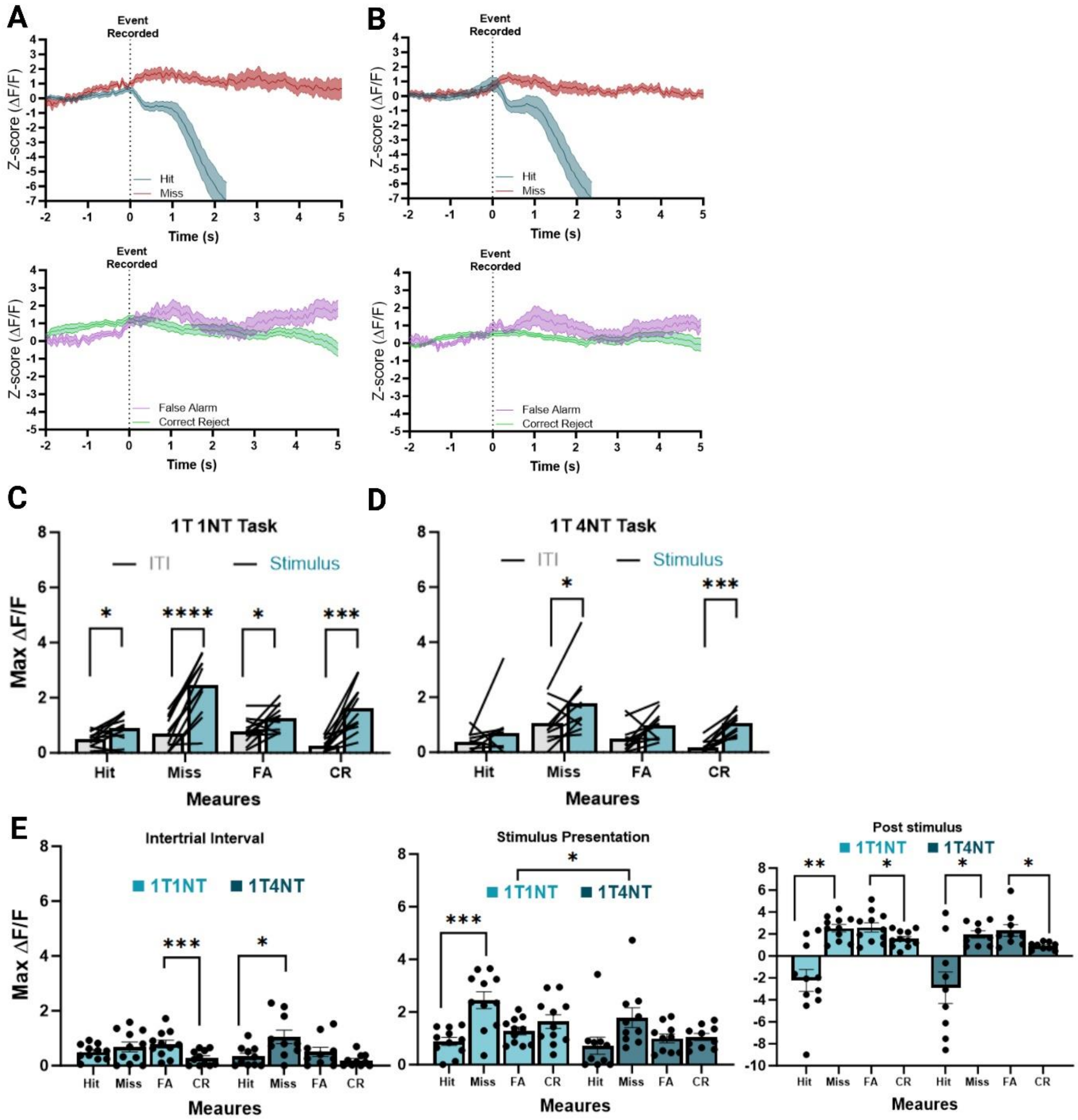


**Supplementary Figure 1. Additional measures of prefrontal PVN activity during the 1NT and 4NT tasks.**

**A.** Averaged signal traces from prefrontal PVNs aligned to the time of responses being recorded on the 1NT. Responses to target stimuli are shown on top and responses to non-target stimuli are shown below.

**B.** Averaged signal traces from prefrontal PVNs aligned to the time of responses being recorded on the 1NT. Responses to target stimuli are shown on top and responses to non-target stimuli are shown below.

**C.** The maximum ΔF/F comparing the intertrial interval (ITI) and stimulus presentation for each measure on the 1NT task. There was a significant effect of task period ($F_{1, 10} = 103.3$, $p < 0.0001$), with significantly higher PVN activity peaks during stimulus presentation on a Hit ($p = 0.0425$), Miss ($p < 0.0001$), False Alarm ($p = 0.0425$), and Correct Rejection ($p = 0.0013$).

**D.** The maximum ΔF/F comparing the intertrial interval (ITI) and stimulus presentation for each measure on the 4NT task. There was a significant effect of task period ($F_{1, 9} = 25.95$, $p < 0.0007$), with significantly higher PVN activity peaks during stimulus presentation on a Miss ($p = 0.0160$) and Correct Rejection ($p = 0.0065$), but not a Hit ($p = 0.1981$) or False Alarm ($p = 0.1075$). n = 11 PV:Cre mice.

Error bars indicate mean ± SEM. **** $p < 0.0001$, *** $p < 0.001$, ** $p < 0.01$, * $p < 0.05$.

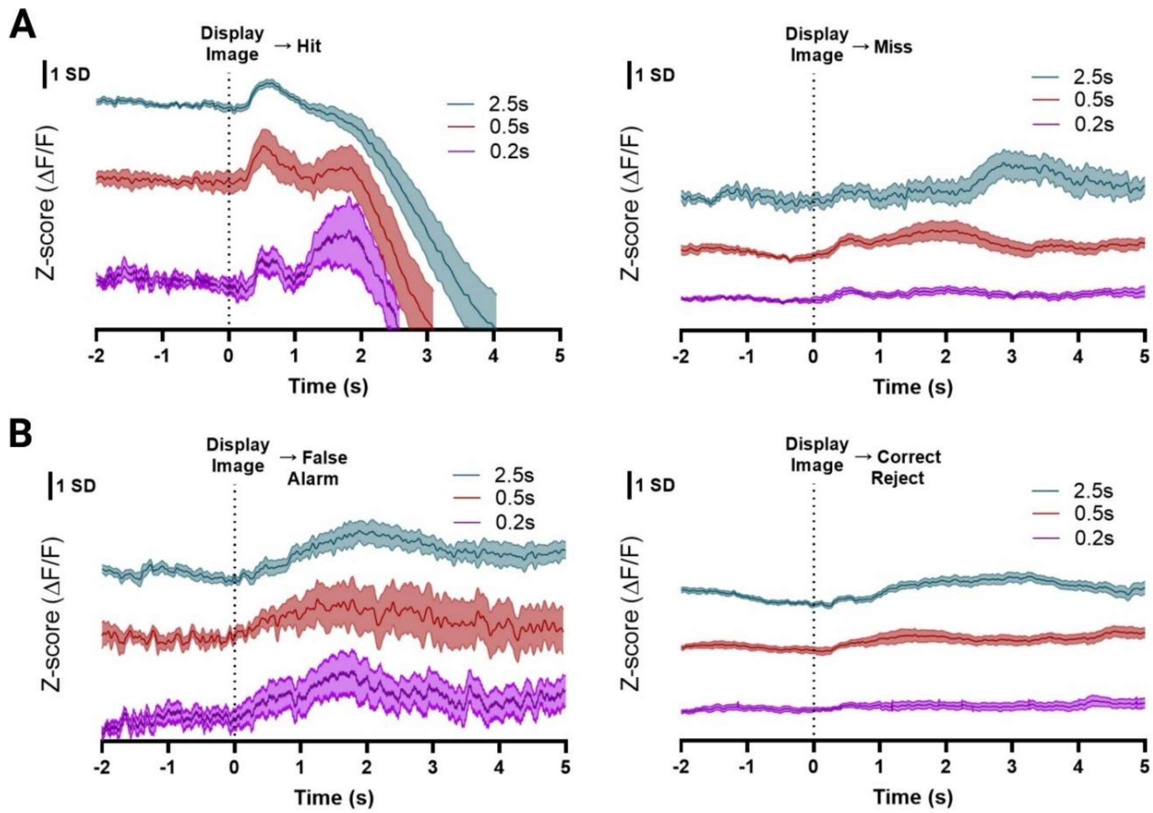


**Supplementary Figure 2. Comparison of prefrontal parvalbumin activity during response types across sessions with reduced stimulus presentation durations.**

**A.** Averaged signal traces from prefrontal PVNs during target presentations across sessions with different stimulus presentation duration. Traces are shown for Hit (left) and Miss responses (right). Baseline session (2.5s) is shown in teal, moderate difficulty session (0.5s) is shown in red, and high difficulty session (0.2s) is shown in purple.

**B.** Averaged signal traces from prefrontal PVNs during non-target presentations across sessions with different stimulus presentation duration. Traces are shown for False Alarm (left) and Correct Reject responses (right). Baseline session (2.5s) is shown in teal, moderate difficulty session (0.5s) is shown in red, and high difficulty session (0.2s) is shown in purple. Scale bars are 1 standard deviation (SD). n = 9 PV:Cre mice.

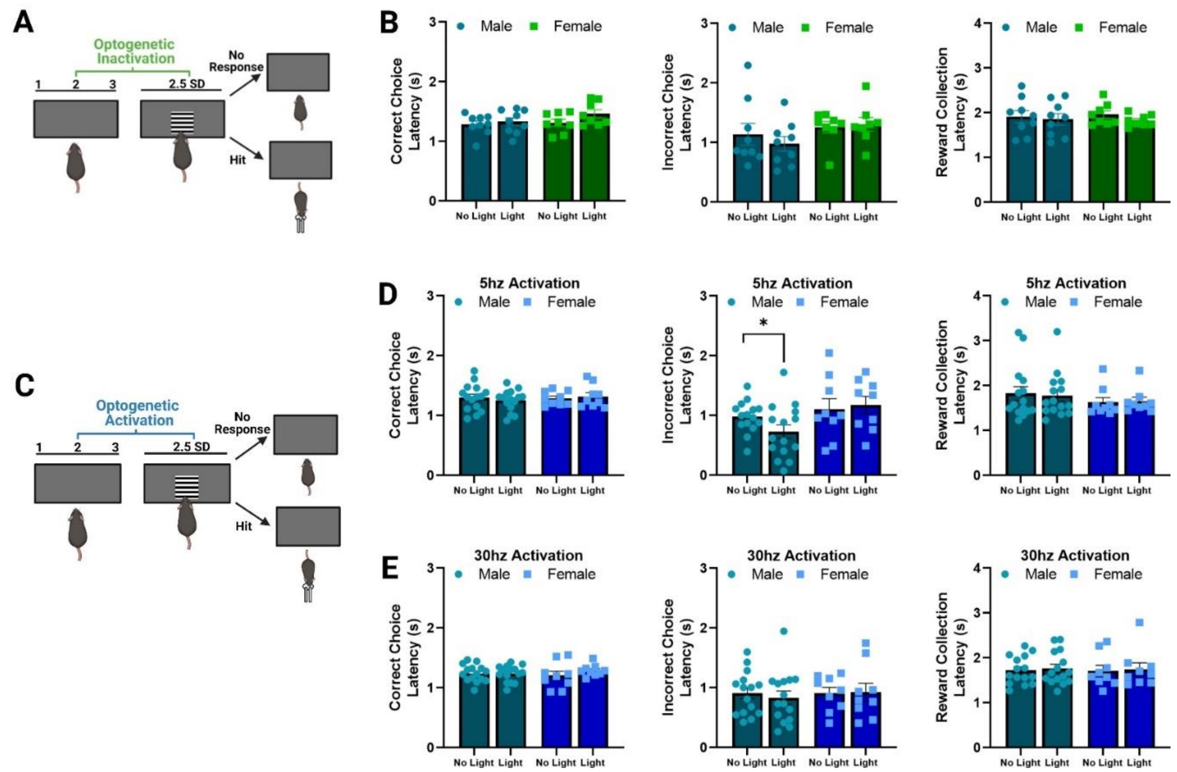


**Supplementary Figure 3. Effects of optogenetic inhibition and frequency-specific activation of prefrontal parvalbumin neurons on latency measures during the 1NT task.**

**A.** PVNs in prefrontal cortex were optogenetically inhibited during the 1NT task.

**B.** There was no effect of PVN inhibition across any latency measure. Measures included time (seconds) to make a hit response (correct choice latency), time to make a false alarm (incorrect choice latency), and time to collect reward following a hit response (reward collection latency. N = 9 male, 8 female mice.

**C.** PVNs in prefrontal cortex were optogenetically stimulated during the 1NT task.

**D.** The effects of 5hz stimulation on latency measures. While there was no effect of stimulation on correct choice latency (Two-way RM ANOVA: $F_{1,22} = 0.097$, $p = 0.759$), there was a significant effect of sex with 5hz stimulation for incorrect choice latency (main effect of sex: $F_{1,22} = 4.478$, $p = 0.046$; no main effect of optogenetics or interaction). Reward collection latency was also unaffected. n = 15 male, 9 female mice.

**E.** There were no effects of 30hz stimulation on any latency measures.

Error bars indicate mean ± SEM. * $p < 0.05$.

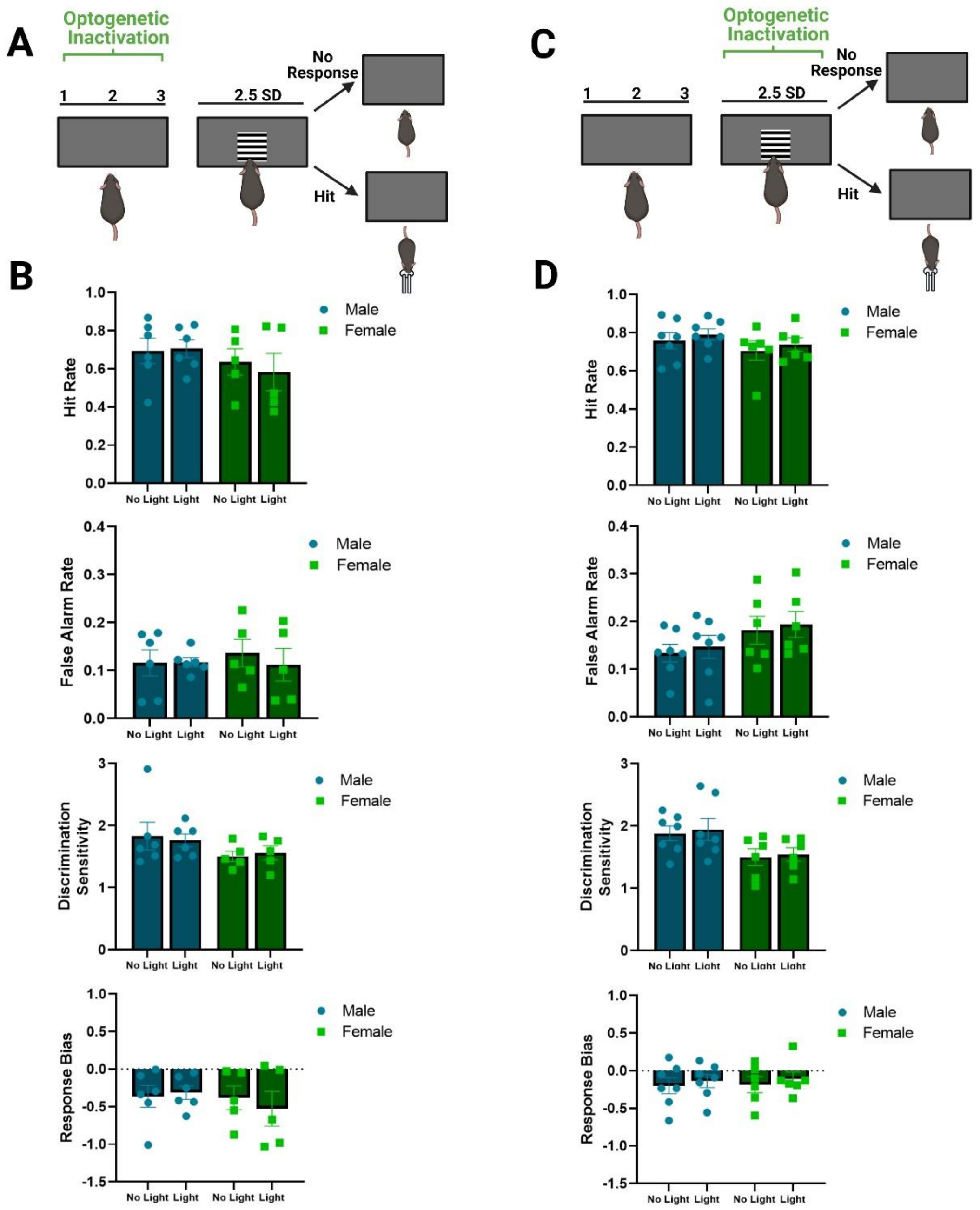


**Supplementary Figure 4. Task-phase specific inhibition of prefrontal parvalbumin neurons during the 1NT task.**

**A.** Intertrial interval (ITI) only optogenetic inhibition of prefrontal PVNs during the 1NT task.

**B.** There was no effect of prefrontal PVN inhibition on any task performance measure during ITI inactivation. n = 6 male and 6 female mice.

**C.** Image presentation only optogenetic inhibition of prefrontal PVNs during the 1NT task.

**D.** There was no effect of prefrontal PVN inhibition on any task performance measure during stimulus inactivation. n = 7 male and 6 female mice.

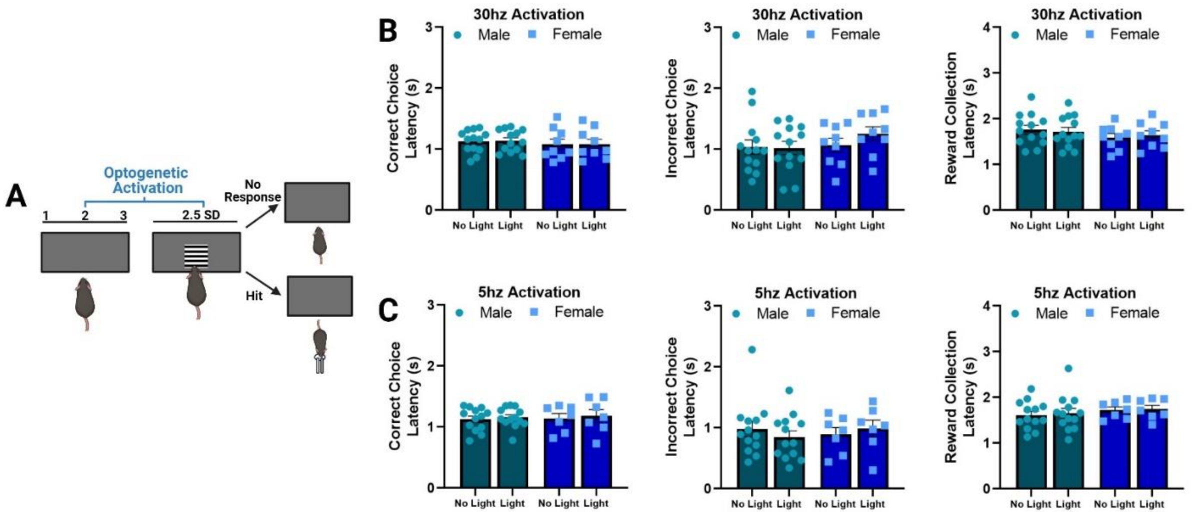


**Supplementary Figure 5. Effects of optogenetic frequency-specific activation of prefrontal parvalbumin neurons on latency measures during the 4NT rCPT.**
**A.** Optogenetic stimulation of prefrontal PVNs during the 4NT rCPT.
**B.** There were no effects of 30hz stimulation on any latency measures (n = 13 males, 9 females).
**C.** There were no effects of 5hz stimulation on any latency measures (n = 13 males, 7 females).

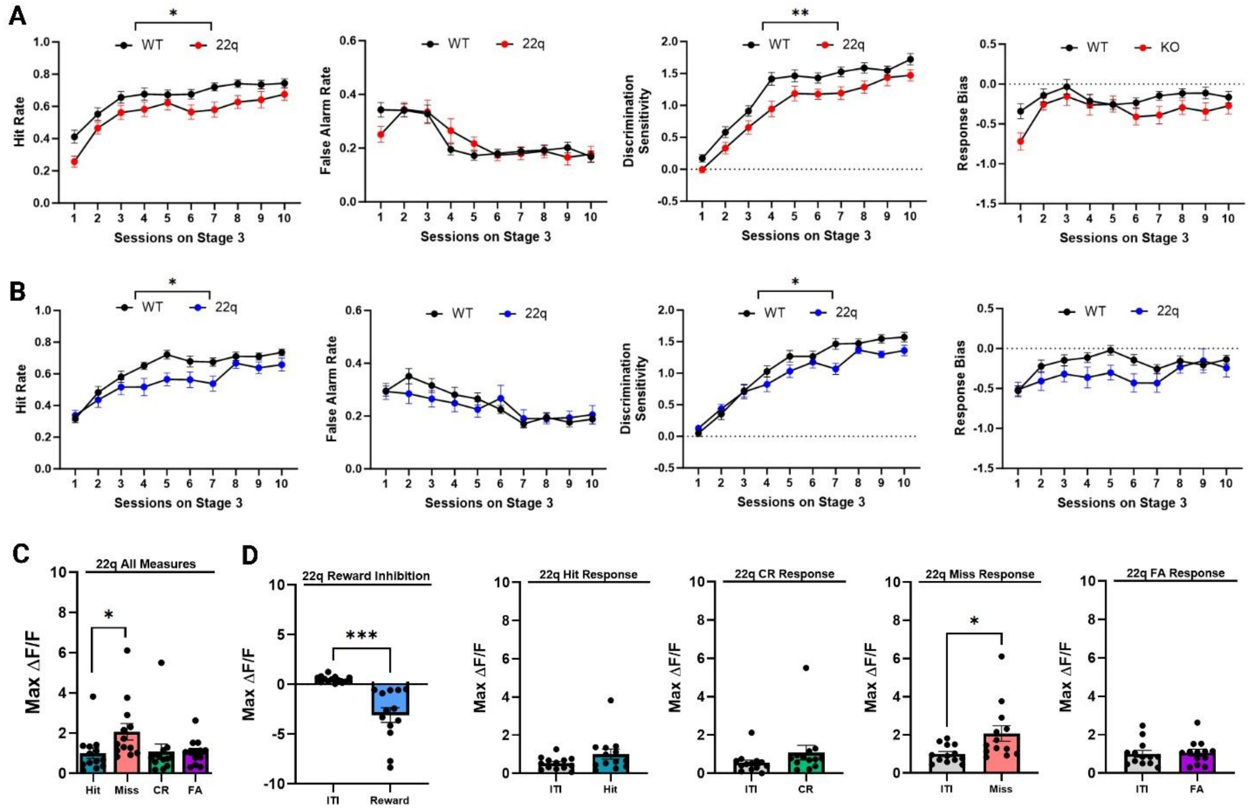


**Supplementary Figure 6. Characterization of the attention deficit profile in male and female 22q mice on the rodent continuous performance task**.
**A.** Male 22q mice (n = 16) display a significant impairment in target detection and discrimination compared to wild-type controls (n = 20) across 1NT task acquisition. One-way RM ANOVA

showed a significant reduction in hit rate ($F_{1,34}$ = 6.759, p = 0.014), and discrimination sensitivity ($F_{1,34}$ = 11.671, p = 0.002), but no effect on false alarm rate ($F_{1,34}$ = 0.003, p = 0.958) or response bias ($F_{1,34}$ = 3.106, p = 0.087). There was a significant effect of session for all measures independent of an interaction with genotype (p < 0.001).

**B.** Female 22q mice (n = 16) display a significant impairment in target detection and discrimination compared to wild-type controls (n = 19) across 1NT task acquisition. One-way RM ANOVA showed a significant reduction in hit rate ($F_{1,33}$ = 5.591, p = 0.021), and a genotype x session interaction for discrimination sensitivity ($F_{9,297}$ = 2.691, p = 0.005), but no effect on false alarm rate ($F_{1,33}$ = 0.1761, p = 0.6775) or response bias ($F_{1,33}$ = 3.521, p = 0.069). There was a significant effect of session for all measures independent of an interaction with genotype (p < 0.001).

**C.** Peak ΔF/F from PVN signals during the stimulus presentation period for each response type in 22q mice. PVN activity significantly differed across response types (One-way RM ANOVA: $F_{2.24, 24.60}$ = 4.272, p = 0.020). This was specific to responses to target images, where PVN activity was significantly elevated during Miss responses compared to Hits (paired t-test: $t_{12}$ = 2.792, p = 0.016).

**D.** Peak ΔF/F from PVN signals during stimulus presentation compared to the ITI for all task measures in 22 mice. 22q mice display significant suppression of prefrontal PVN activity when reward is collected following a correct Hit (paired t-test: $t_{12}$ = 4.746, p = 0.0004) and significant recruitment of PVN activity during Miss responses to a target (paired t-test: $t_{12}$ = 2.849, p = 0.015. No additional measures were associated with significant changes in PVN activity.

Error bars indicate mean ± SEM. *** p < 0.001, ** p < 0.01, * p ≤ 0.05.

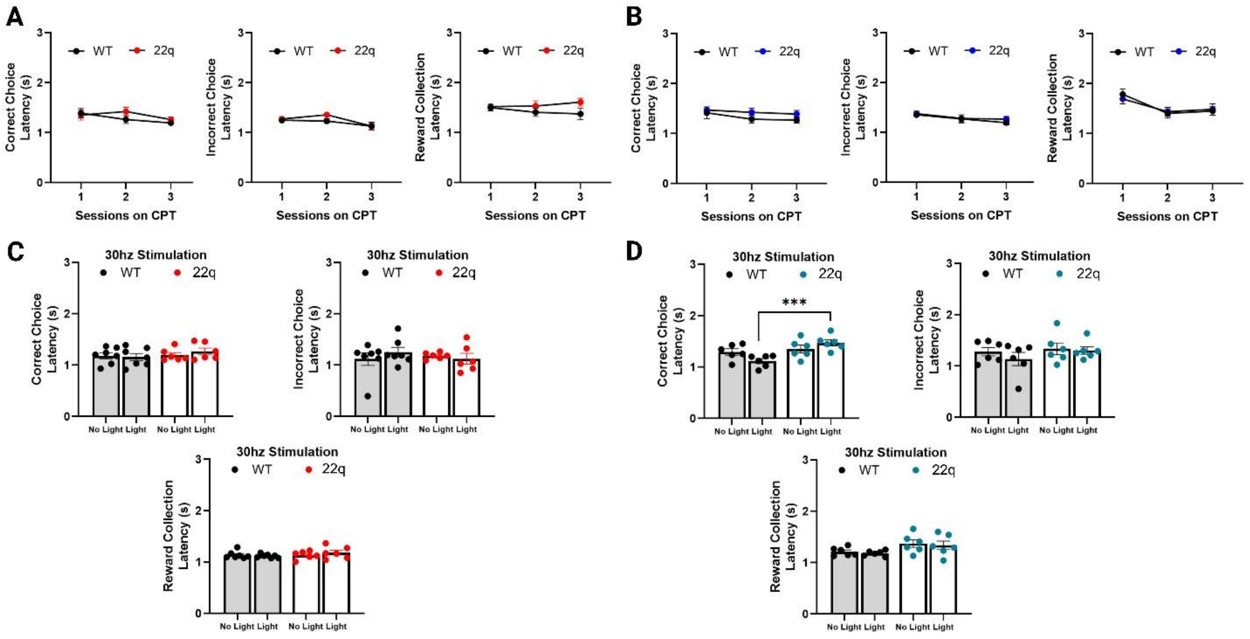


**Supplementary Figure 7. Latency measures during 1NT training and 30hz stimulation of prefrontal parvalbumin neurons in 22q mice and wild-type controls.**

**A-B.** There were no significant differences on any latency measures between 22q mice and wild-type controls during the first 3 training sessions on the 1NT task. (Left) Males are shown in red (n = 7 wild-types and 6 22q mice), and (right) females are shown in blue (n = 6 per group).

**C.** There were no effects of optogenetic stimulation or genotype on any latencies in male mice.
**D.** There was a transient interaction between genotype and optogenetic stimulation for correct choice latency in female mice (Two-way RM ANOVA: $F_{1,10} = 8.276$, $p = 0.016$; main effect of genotype: $F_{1,10} = 9.076$, $p = 0.013$) during the light condition (Post hoc unpaired t-test: $t_{10} = 4.719$, $p < 0.001$). There were no effects of genotype or optogenetic stimulation on other measures. n = 6 per group.

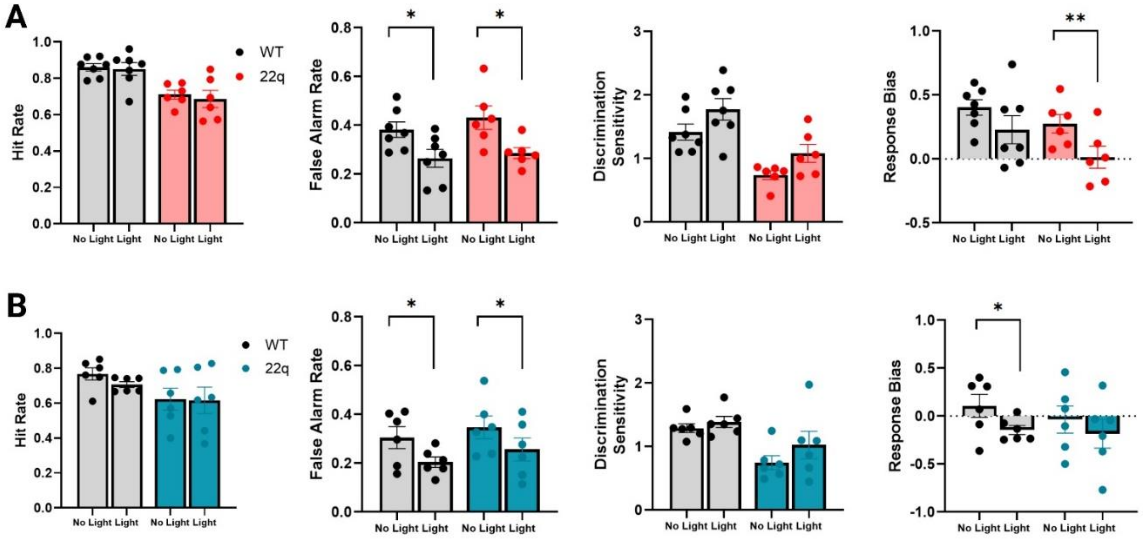


**Supplementary Figure 8.** Within-subjects effects of 30hz stimulation on prefrontal parvalbumin neurons during attention in 22q mice and wild-type controls.
**A.** Stimulating prefrontal PVNs significantly reduced false alarm rate in both male 22q (red, paired t-test: $t_5 = 3.728$, $p = 0.014$) and wild-type controls (paired t-test: $t_6 = 3.186$, $p = 0.019$). Additionally, 30hz stimulation significantly reduced response bias (Two-way RM ANOVA: $F_{1, 11} = 14.684$, $p = 0.003$) specifically in 22q mice (paired t-test: $t_5 = 5.827$, $p = 0.002$) but not wild-types (paired t-test: $t_6 = 2.190$, $p = 0.071$). n = 7 wild-type and 6 22q mice.
**B.** Stimulating prefrontal PVNs significantly reduced false alarm rate in both female 22q (blue, paired t-test: $t_5 = 3.711$, $p = 0.013$) and wild-type controls (paired t-test: $t = 2.716$, $p = 0.042$). n = 6 per group. Additionally, 30hz stimulation significantly reduced response bias (Two-way RM ANOVA: $F_{1, 10} = 14.14$, $p = 0.004$) specifically in wild-type mice (paired t-test: $t_5 = 3.363$, $p = 0.031$) but not 22q mice (paired t-test: $t_5 = 2.508$, $p = 0.054$).

Error bars indicate mean ± SEM. ** $p < 0.01$, * $p < 0.05$.

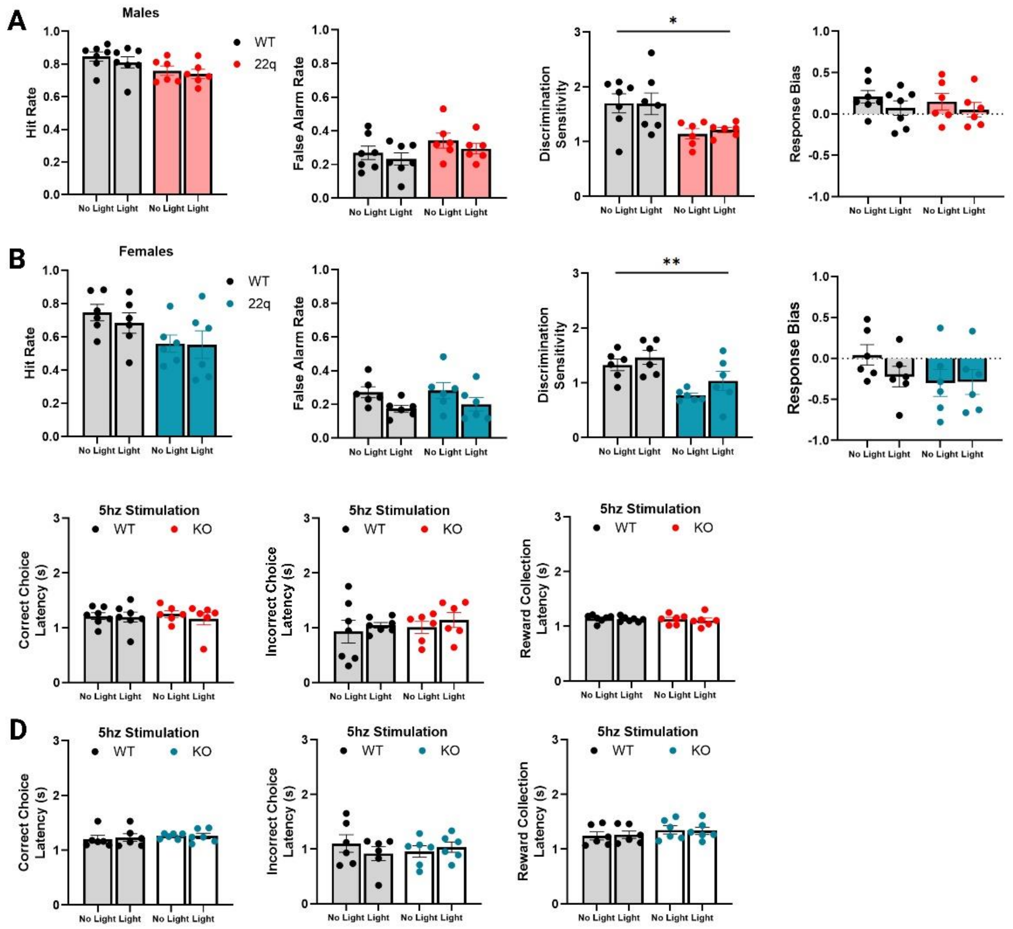


**Supplementary Figure 9.** 5hz stimulation of prefrontal parvalbumin neurons had no effect on discrimination sensitivity in 22q mice. **A.** The effects of stimulating prefrontal PVNs at 5hz in male 22q and wild-type control mice during the 1T, 1NT go/no task. 22q mice displayed significant reduced discrimination sensitivity compared to wild-type controls (Two-way ANOVA: $F_{1, 11} = 7.937$, $p = 0.017$), and performance between groups did not differ across any other measure. Further, there was no effects of optogenetic stimulation ($F_{1,11} = 0.076$, $p = 0.788$), showing that stimulating PVNs at 5hz had no effect on task performance. n = 7 wild-type controls and 6 22q mice.

**B.** The effects stimulating prefrontal PVNs at 5hz in female 22q and wild-type control mice during the 1T, 1NT go/no task. 22q mice displayed significantly reduced discrimination sensitivity compared to wild-type controls (Two-way ANOVA: $F_{1, 10} = 11.956$, $p = 0.006$), and performance between groups did not differ across any other measure. There were no significant effects of optogenetic stimulation ($F_{1,10} = 4.627$, $p = 0.057$), showing that stimulating PVNs at 5hz was unable to rescue deficits in task performance. n = 6 mice per group.

**C.** There were no effects of 5hz stimulation on task latency measures in (C) male mice or

**D.** female mice in either group.

Error bars indicate mean ± SEM. ** $p < 0.01$, * $p < 0.05$.
